# Trajectory fit alone does not identify causal edges: a white-box masked ODE benchmark for trans-omics digital twins of drug action mechanisms

**DOI:** 10.64898/2026.09.22.753490

**Authors:** Zhulv Zhang, Lirong Jia, Yanli Pan, Haiyu Zhao

**Author notes:** Correspondence: Yanli Pan; Haiyu Zhao.

## Abstract

Dynamic multi-omics models infer drug mechanisms from trajectories, but trajectory accuracy alone cannot verify that learned regulatory edges are mechanistically correct. We asked to what extent fitting time-series dynamics identifies intervenable causal edges across observation and optimization budgets. We built white-box ordinary differential equations with candidate-edge masks for gene-expression, protein, signalling and metabolite dynamics, using a mouse-liver trans-omics topology and ground-truth semi-synthetic time series. Under noise-free, complete, densely sampled data, from-scratch training ranked edges near chance (edge-class area under the receiver-operating characteristic curve, 0.48–0.53); the unattainable oracle-initialized upper bound reached 0.94–0.99 on the same metric. Unobserved layers absorbed error. In ground-truth knockout tests, development-batch endpoint-sign accuracy was 67.9% for one-hop non-convergent targets but 33.3% for two-hop targets. Across tested practical budgets, fitting did not yield reliable edge recovery; trajectory fit alone therefore does not establish mechanistic identification. This benchmark supports mechanistic audits and exploratory per-prediction confidence grading in multi-omics models.

## 1. Introduction

Drug action is dynamic and multi-layered. A stimulus such as insulin propagates from signalling through transcription and protein abundance to metabolism on timescales from minutes to hours [1–3]. Understanding a drug’s mechanism of action increasingly means predicting these molecular trajectories rather than cataloguing static associations. This is the promise of molecular digital twins, computational replicas that can be perturbed in silico before any experiment is run [4,5]. This capability is especially relevant for multi-component interventions, including traditional Chinese medicine (TCM) formulas, which act on multiple targets and can produce responses across multiple omic layers. Their mechanistic interpretation remains an open problem despite intensive network-pharmacology efforts [6,7].

Trainable dynamic models fitted to time-series multi-omics provide a route to predicting these responses. ODE-constrained learning, universal differential equations and knowledge-guided machine learning [8–10] provide general modeling frameworks, but they do not remove the validation problem: for any such model, including the white-box ODE we build below, fitting observed trajectories does not by itself establish that the learned regulatory edges are mechanistically correct. For static network inference the community learned this lesson through two decades of ground-truth benchmarking: the DREAM challenges established that accuracy claims are meaningless without blinded gold standards [11–14]. For dynamic, multi-layer, genome-scale models, no equivalent ground truth has existed, and the question “to what extent does fitting time-series dynamics identify intervenable causal edges across observation and optimization budgets?” has remained untestable rather than untested. Recent perturbation-prediction benchmarks offer a related caution from the application side: deep models trained on observational data do not yet outperform simple linear baselines in predicting unseen perturbation effects [15]. Although that task differs from dynamic edge identification, it reinforces the need for evaluation that directly tests a model’s claimed capability. The stakes are now widely recognized: identifiability and perturbation generalization have been named core challenges for biomedical modeling [16], and “Challenge 1: Regulatory and signaling interactions” heads the fifteen challenges recently codified for generative AI in cell biology [17]. An empirical identifiability-budget benchmark oriented to mechanistic computation is therefore needed.

Existing identifiability theory does not close this gap at the relevant scale. Structural and practical identifiability analysis [18–21] certifies whether parameters are identifiable in principle or locally at a fitted point, typically for systems with tens of variables. Recent work on hybrid neural ODEs applies Hessian-based local analysis and reports qualitative compensation between neural and mechanistic terms as a source of non-identifiability [22]. That analysis is local and restricted to small systems. By contrast, the system built in this paper has 2,106 molecular species and ∼360,000 candidate edges (4,912 realized in the semi-synthetic ground-truth network); at that scale, Hessian-based local analysis is computationally infeasible. Binary identifiability certificates also fail the practitioner’s real questions: how far is failure, what caused it, and what would fix it?

Here we turn identifiability into an operational, empirically measurable budget. This budget is a component-wise account of how observation conditions, initialization and saturation each contribute to operational edge identifiability, not a single composite score. We build a white-box masked ODE on the real trans-omics topology of insulin action in mouse liver (GSE166336). Every term is mechanistic (production, first-order decay, and tanh regulation); prior knowledge enters only as edge masks that define the candidate set. A semi-synthetic engine on the same topology provides ground-truth networks with realistic noise, missingness and sampling grids, in the spirit of GeneNetWeaver [13] but dynamic and trans-omic. We call the fully known semi-synthetic system produced by this engine the ground-truth universe and refer to the engine as the ground-truth generator. Throughout, edge-class AUROC denotes AUROC computed separately within each of the six edge-class candidate domains (GE, SG, EC, MM, FB and PS), not an AUROC pooled across edge classes. Four instruments then measure where identification fails. (i) An oracle-init basin curve tracks edge-class AUROC (area under the receiver-operating characteristic curve) as a function of initialization noise around the ground truth. (ii) A held-out-layer corruption assay isolates gradient-mediated contamination with freeze probes. (iii) A tanh-saturation audit quantifies numerically silent regulator columns. (iv) An ideal-budget ceiling test separates data limits from algorithmic limits.

Five findings emerge. First, a static finite-difference (FD) + least absolute shrinkage and selection operator (LASSO) baseline performs at chance, illustrating quasi-steady-state blindness and motivating dynamic fitting. Second, edge recovery is strongly initialization-dependent: from-scratch training remains at chance even with noise-free, fully observed data with 49 time points at 5-min spacing (edge-class AUROC 0.48–0.53), whereas the unattainable oracle-initialized ideal-data upper bound reaches 0.94–0.99 on the same metric. Third, held-out layers act as corruption sinks; intervention dissection identifies the bias channel as the largest compensation carrier and the P-pathway kinetic block as a secondary carrier.

Fourth, tanh saturation can suppress regulator-column gradients at realistic operating-point offsets. Fifth, a 12-knockout in-silico validation battery shows that intervention reliability depends on network position: one-hop non-convergent predictions retain trajectory shape and direction (endpoint sign 67.9%), whereas two-hop predictions are not reportable (endpoint sign 33.3%). Across these findings, the four instruments distinguish failures of initialization, observation, saturation and data budget, motivating per-prediction confidence triage rather than a global accuracy score.

These findings lead to four practical prescriptions: first, initialization and priors are necessities, not refinements, with the caveat that priors act as candidate-set masks and basin placement, while sign priors alone do not suffice (§2.4); second, any layer carrying a mechanistic claim must be measured, because held-out layers are corruption sinks (here, the held-out P layer failed outright, whereas the held-out S layer remained at or below chance); third, scale-matched initialization plus saturation audits belong in digital-twin reporting standards; and fourth, digital-twin predictions should ship with per-prediction confidence grades (one-hop, non-convergent / directional-only / not reportable), degrading automatically when path distance is unknown; the present grades are a preliminary calibration. We discuss implications for mechanism elucidation of multi-component drugs, including TCM formulas, as the motivating hardest case.

## 2. Results

### 2.1 A white-box masked ODE benchmark on real trans-omics topology (Fig. 1)

We constructed the benchmark on the insulin-stimulated mouse-liver trans-omics dataset of Matsuzaki et al. [23] (transcriptome: GEO GSE166336; proteome: ProteomeXchange PXD022728; phosphoproteome: PXD022823; metabolome: source-publication Tables S2–S3). The retained four-layer state space comprises 691 transcripts (G), 691 proteins (P), 691 phosphosite signals (S) and 33 metabolites (M): 2,106 molecular species in total. Prior knowledge from KEGG (Kyoto Encyclopedia of Genes and Genomes) KGML (KEGG Markup Language) files and curated literature defines candidate edge masks in six classes (gene–gene, GE; signalling-to-gene, SG; enzyme–metabolite, EC; metabolite–metabolite, MM; feedback, FB; protein–phosphosite, PS). The three counts along this chain are distinct: the six masks define ∼360,000 candidate edges (class-wise counts in §4.2; retention probabilities in §4.4); the full GE prior mask (deduplicated pairs; sources in §4.2) contains 3,885 edges; and the semi-synthetic ground-truth network comprises 4,912 edges subsampled within the ∼360,000 candidate masks. Trimming the GE mask to the measured gene list leaves the 3,878 GE candidate edges of the real-data fit (§2.3); the seven removed edges involve genes outside the measured list. For the semi-synthetic benchmark, slicing the full GE prior mask to the KGML-defined subnetwork yields 362 candidate GE edges, of which 115 are realized in the ground-truth network; this candidate domain is distinct from the 3,878-edge GE mask used for the real-data fit. The ODE contains no other terms, so every learned edge is readable and auditable (white-box). Each equation combines masked tanh regulation (except for the proteome layer, which has translation coupling, bias and decay but no tanh term), linear production gated by a softplus-calibrated layer coupling (a smooth positive-part function), and first-order decay; tanh was chosen as the minimal auditable sigmoid (a smooth saturating, Hill-type response).

**Figure 1.**
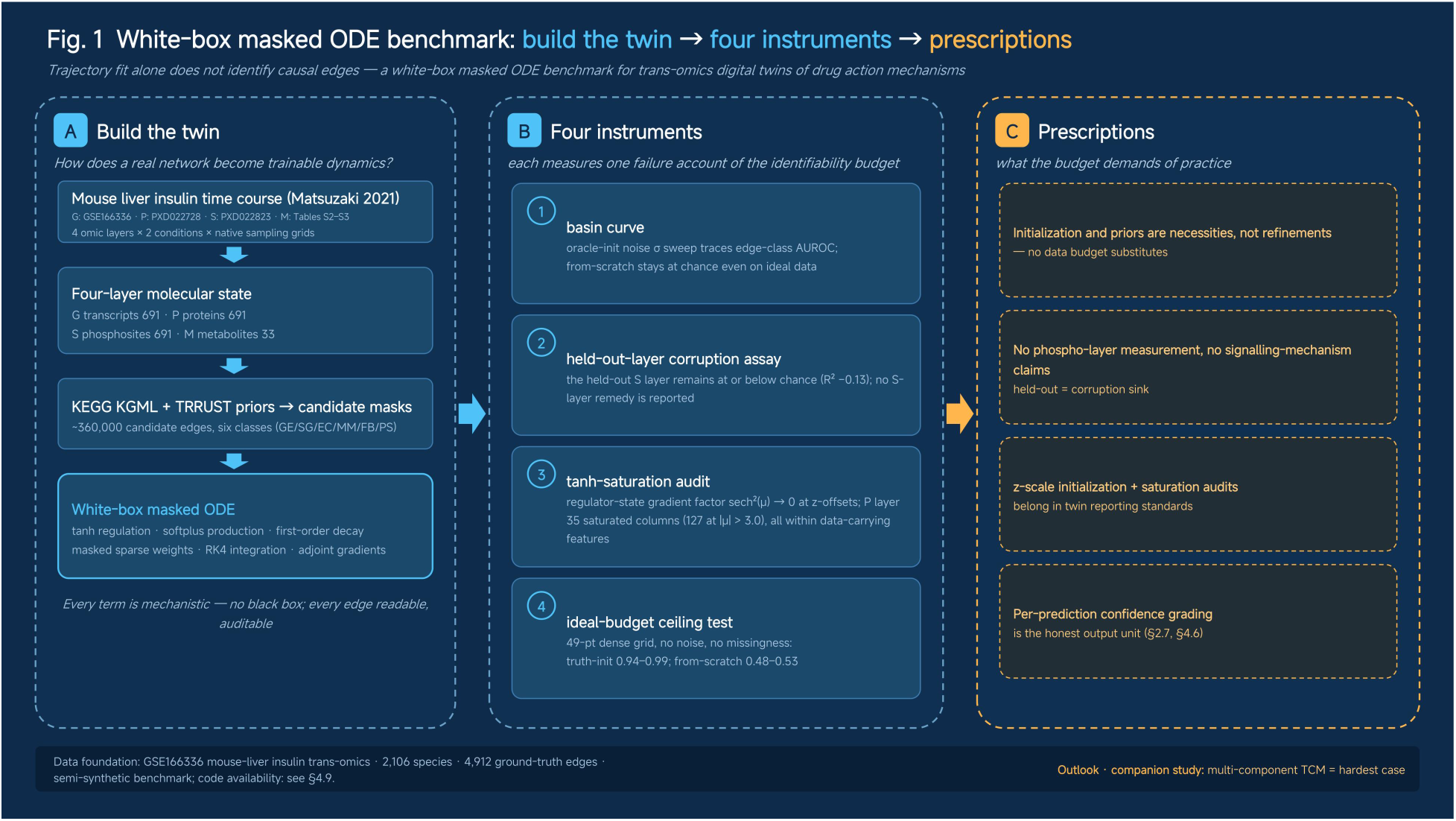
Benchmark framework on the GSE166336 mouse-liver insulin time course: four omic layers under two conditions, candidate-edge prior masks, the white-box masked ODE (A), the four evaluation instruments (B), and the resulting prescriptions (C).

Training uses fourth-order Runge–Kutta (RK4) integration with adjoint gradients (backpropagation through the ODE solver) and, for sparsification, a proximal soft-thresholding step applied after every optimizer update (w ← sign(w)·max(|w| − τ, 0); Methods). A semi-synthetic engine on the identical topology samples ground-truth networks from the candidate masks and generates trajectory pairs (insulin/control) with realistic observation noise (per-replicate σ = 0.1 in z-space; three-replicate averaging gives an effective σ ≈ 0.058), feature-level missingness (P: 76.6%, S: 82.2% of features entirely unobserved, mirroring the real data) and the real sampling grids; an ideal-data mode (49-point dense grids, zero noise, zero missingness) probes the observation-budget ceiling.

All evaluations use the same metric protocol: identical candidate sets, identical flat-prediction baselines (predicting the observation mean), and identical statistics (edge AUROC/AUPRC (area under the precision–recall curve)/Top-k; held-out layer R²) across arms. We chose the insulin-response topology because it is one of the few public systems with four-layer trans-omics time resolution: it supplies the realistic network statistics (sparsity, inter-layer connection density, edge-class composition) that random or scale-free graphs cannot. The benchmark uses the insulin-response topology without attempting a complete biochemical reconstruction. It asks: given this backbone and semi-synthetic data generated on it, to what extent does fitting dynamics recover the backbone’s edges across observation and optimization budgets? The instruments below are dataset-agnostic, and the audited failure modes are attributes of the training problem; we return to their generality in the Discussion. Conclusions are drawn for this model class: a functional-form-mismatch audit within the Hill family (§2.8) supports rank-level recovery but does not establish these accounts beyond the tested functional forms.

### 2.2 A static FD + LASSO baseline is at chance: quasi-steady-state blindness (Fig. 2C)

Under the same data (semi-synthetic universe), candidate sets and metrics, a finite-difference + per-target LASSO baseline, the static workhorse of network inference [24], performs at chance on five of six edge classes, each scored within their respective candidate domains (GE slightly above: 0.567; AUROC: SG 0.496, PS 0.500, FB 0.502, EC 0.500, MM 0.497). Its PS Top-k precision, the proportion of true edges among the k highest-scored candidates, is 0.995% at a 1.0% candidate base rate (k = 3,509; including all candidates tied at the kth score selected the full candidate set). The ODE trained from a mildly perturbed oracle (σ = 0.1) reaches 0.805–0.859. This is consistent with a structural quasi-steady-state blindness: at realistic sampling, most molecular states are barely moving, so the finite-difference derivative carries the source’s rate of change, not its level. Static regression is thus blind to exactly the signal that dynamic fitting exploits. Combined with class-dependent predictor dimensionality (2–594 candidates per target) and the absence of cross-node coupling constraints, chance-level performance is the expected outcome, echoing two decades of DREAM experience [11,14]. As an independent method control, dynGENIE3 attained a tie-averaged (§4.6) AUROC of 0.607 ± 0.016 (mean ± half-width of the 95% t-interval across the five seeds, Table S4) on the ideal-data mode at the 1000-tree setting and 0.562 (mean across the same five seeds) on the realistic-observation mode (§4.7, Table S4; two observation modes over the same ground-truth universe, §4.4); scoring was confined to the GE candidate domain because per-edge scoring across the full 2,106-node universe is computationally impractical for tree-ensemble methods (§4.7).

**Figure 2.**
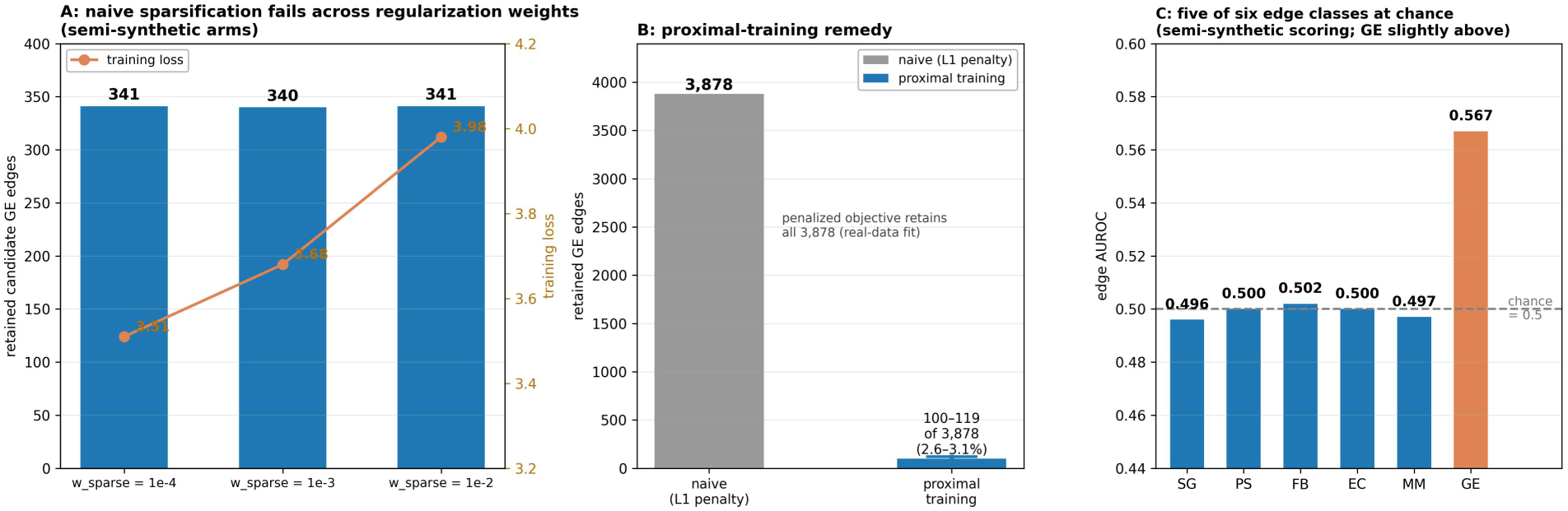
A: naive regularization-weight sparsification failure modes (semi-synthetic universe). B: proximal training on the real-data fit — 100–119 of 3,878 candidate GE edges retained, versus all 3,878 under the penalized objective. C: static LASSO baselines remain near chance across edge classes (GE slightly above at 0.567).

### 2.3 Naive sparsification fails; proximal training restores sparsity on the real-data fit (Fig. 2)

Standard practice, Adam with an L1 penalty, does not sparsify this model. It retains 341 of the 362 candidate GE edges in the semi-synthetic universe, 94% of the candidate set, effectively no sparsification, because oscillating gradients never land exactly on zero. On the real-data fit, adding a proximal soft-thresholding step after each optimizer update produces a sparse wiring diagram: 100–119 of the 3,878 candidate GE edges are retained across two thresholds (τ = 3×10⁻⁴ and 10⁻³), compared with all 3,878 under the penalized objective alone (Methods). In the semi-synthetic universe, increasing the penalty across two orders of magnitude (w_sparse ∈ {1e-4, 1e-3, 1e-2}, from-scratch arms) does not rescue the failure: retained GE edge counts stay flat (341/340/341). The failure is structural, not a tuning artifact within this range (Fig. 2A).

### 2.4 Initialization dependence and the observation-budget ceiling (Fig. 3)

The basin scan tests how edge recovery changes as initialization moves away from the ground truth. Sweeping oracle initialization noise σ ∈ {0, 0.1, 0.2, 0.3, 0.4} × signal std traces the basin curve: edge-class AUROC degrades from 0.886–0.973 (σ = 0) through 0.805–0.859 (σ = 0.1) to 0.591–0.670 (σ = 0.4). Three cold-start families then probe the realistic condition: from-scratch zero initialization (edge-class AUROC 0.409–0.508), oracle-blind random initialization (Gaussian draws internal to the candidate mask, i.e., no oracle information; two independent training runs, blind3 and blind4; edge-class AUROC 0.46–0.58) and a prior-signed initialization (literature-derived edge signs where available, random signs elsewhere; two independent training runs, prior3 and prior4; edge-class AUROC 0.39–0.57). The cold-start family spans five statistically independent training runs: the zero-initialization arm, re-executed twice (seeds 43/44) with bit-identical results (total loss 8.0000 → 3.5108 over 300 epochs; data term 3.4930; identical learned edge tables; mean edge-class AUROC 0.473) and counted once; the remaining four independent runs come from the other two families. All five statistically independent runs completed training healthily (data-term loss 8.0 → 3.327–3.493), yet every one remains essentially at chance. The weak GE-layer signal has point estimates consistently above chance across all four random/prior arms (AUROC 0.54–0.58; Top-k precision 35–37% vs 31.8% candidate base rate). The failure is unambiguous. *These arms fit the trajectories but do not recover the wiring*: training-loss convergence and edge recovery are separable, and the binding constraint is where optimization starts, not how well it fits.

**Figure 3.**
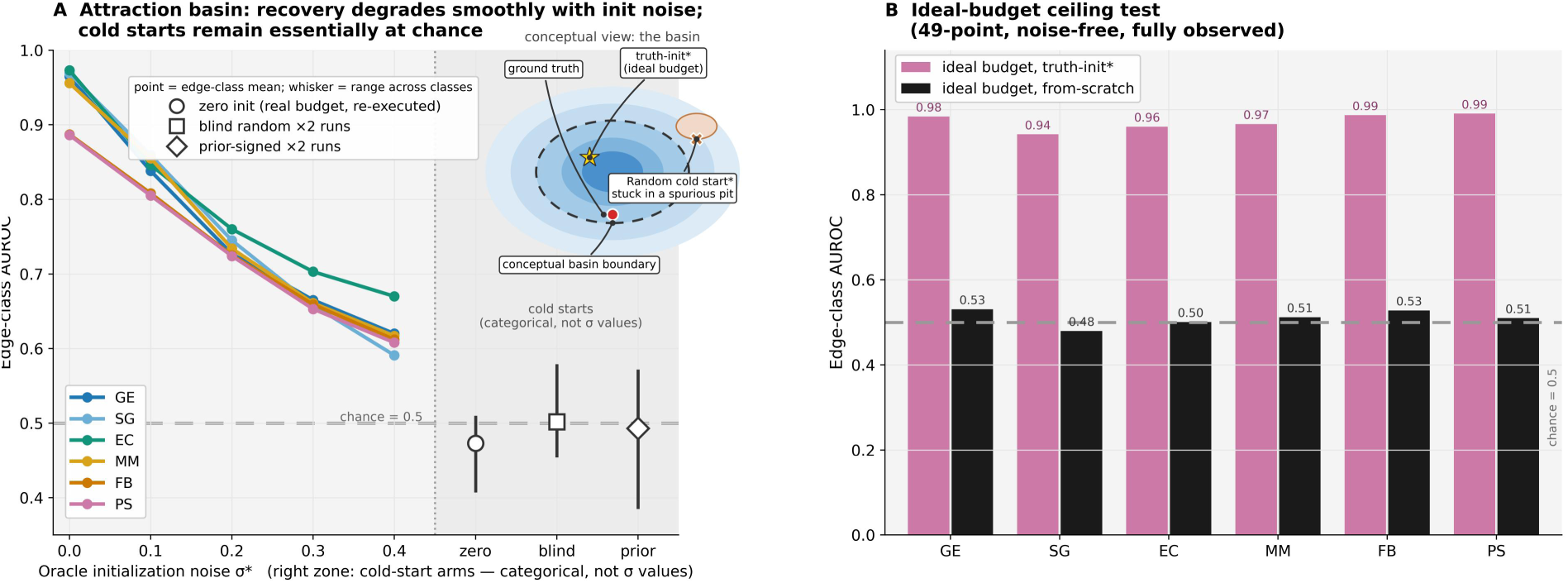
Cold-start arms remain essentially at chance. Oracle-init conditions are upper-bound ablations, not practically attainable. Cold-start arms: the names blind3/blind4 (random init) and prior3/prior4 (prior-signed init) are run IDs of four independent training runs, in addition to the zero-init arm (counted once). Oracle-init noise σ is expressed in units of signal standard deviation. The zero-init arm is deterministic; re-executions are bit-identical and it counts as a single run. * Label marker only; not statistical significance.

Two ideal-budget arms then separate the failure accounts. With 49-point dense, noise-free, fully observed data, the oracle-initialized arm (σ = 0) reaches edge-class AUROCs of 0.942–0.991 across the six classes: most of the difference from the realistic-budget σ = 0 arm is attributable to the combined observation budget—noise, missingness, and sampling density—rather than to the algorithm (the residual gap to 1.0 is attributable to regularization shrinkage (§2.3) and saturation (§2.6)). The from-scratch arm, however, remains at chance even on this ideal data (edge-class AUROC 0.48–0.53, Top-k 4.6% pooled across classes, 1.0–36.5% by edge class). Jointly relaxing these observation-budget constraints does not rescue cold starts; initialization/priors remain the binding constraint under the tested model and optimization protocol. Its at-chance recovery is reproducible across deterministic runs, but this does not distinguish landscape-intrinsic failure from optimizer-specific failure. Quantitatively, the cold-start arms yield numerically lower edge recovery than the σ = 0.4 oracle-init arm. (Initialization details are provided in §4.5.)

### 2.5 Held-out layers are corruption sinks: an intervention-based carrier decomposition (Fig. 4)

Real studies routinely leave entire layers unmeasured. We therefore held out the P and S layers at training time and evaluated the trained model on the withheld trajectories. From-scratch training provides the no-observation reference: without observation, dynamics cannot be invented (held-out R²: P −5.85; S at flat-prediction level; negative values indicate a fit worse than the flat observation-mean baseline). Oracle initialization can fall below this reference because gradient contamination can propagate into incorrect held-out dynamics, whereas from-scratch stays closer to the flat mean.

**Figure 4.**
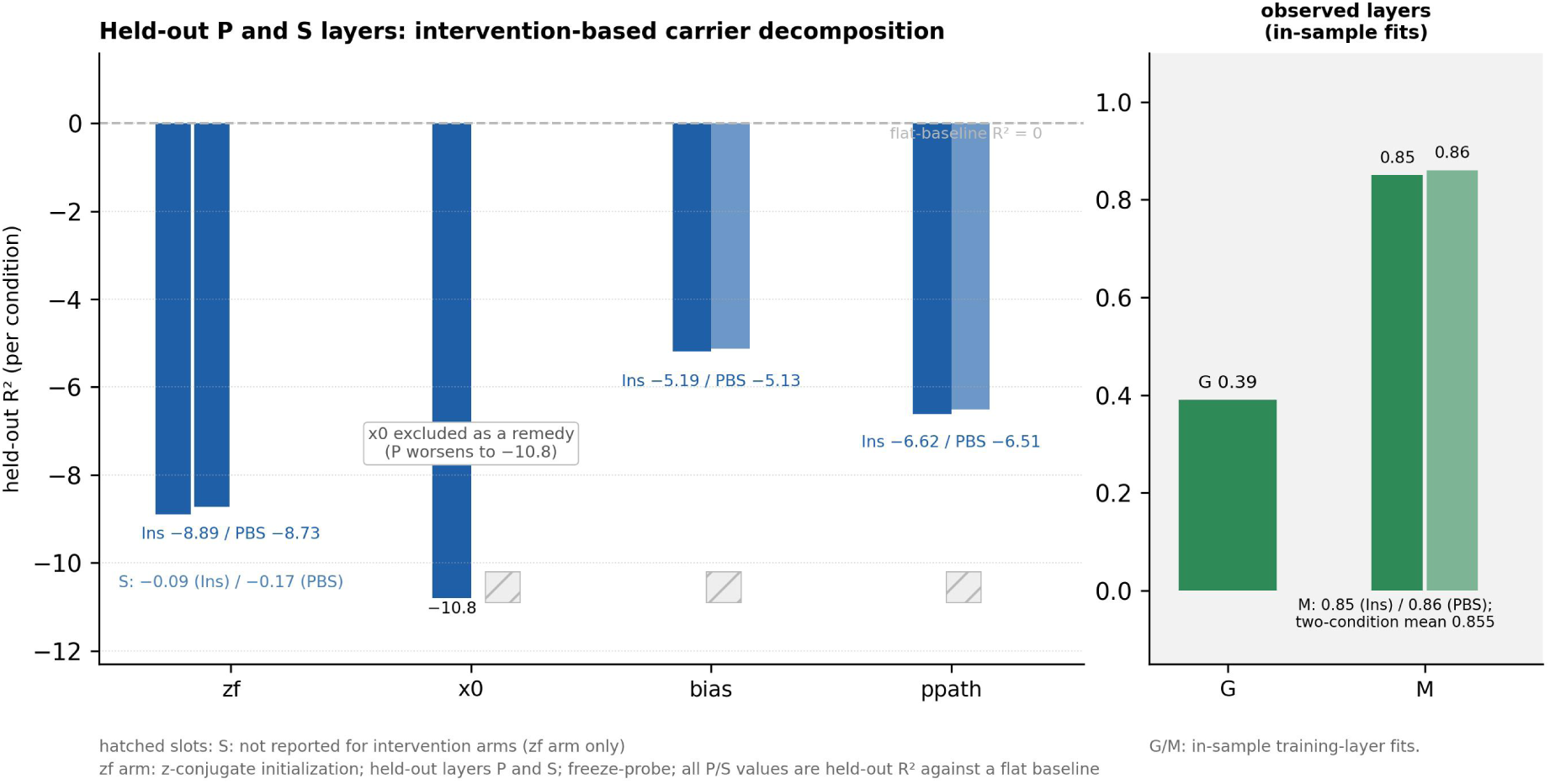
Held-out layers act as corruption sinks. Arm ‘zf’: z-conjugate initialization; held-out layers: P and S; frozen blocks: feedback and phospho-signalling (freeze-probe). The held-out phospho-signalling layer remained at or below chance, and the held-out proteome layer did not recover. Interventional probes identify the bias channel (the per-species additive constant term; §4.3) as the principal compensation carrier of held-out-layer contamination, with the P-pathway kinetic block as a secondary carrier (numerical values in §2.5 and Table S1). G/M values are in-sample training-layer fits; the x0 probe is excluded as a remedy.

Oracle initialization does not rescue held-out layers either. In the z-conjugate frozen-block arm (zf — a freeze-probe that holds selected parameter blocks fixed to locate where the contamination is carried, using z-conjugate initialization, a first-order transform that maps the ground-truth dynamics into the twin’s z-scored coordinates (§4.5), with P and S held out and the feedback and phospho-signalling blocks frozen), the held-out proteome layer fails outright (R² ≈ −8.8; insulin/PBS (phosphate-buffered saline) −8.89/−8.73), and the held-out phospho-signalling layer remains at or below chance (R² −0.13, two-condition mean; per-condition −0.09/−0.17). The observed layers fit normally (G 0.39; M 0.85–0.86; these are in-sample fits and are reported as such).

A four-arm intervention dissection (Table S1) localizes where the failure is carried. Freezing the learnable initial state (x0 probe) makes the P layer worse (−10.8) and is excluded as a remedy. Pinning (fixing a parameter block at its initial value during training) the bias channel raises the P-layer R² from −8.8 to −5.2 (−5.19/−5.13) — the largest recovery, with the other observed layers and the readouts of the other intervention arms unmoved. This identifies the bias channel as the principal compensation carrier (the channel through which the failure signal is carried) of held-out-layer contamination. Pinning the P-pathway coefficients (the P-layer kinetic block: translation coupling, bias and decay) raises R² to −6.6 (−6.62/−6.51), a secondary share. The observed-layer G/M values support that the recovery is intervention-specific rather than a global refit.

The prescription is unchanged and sharp: conclusions about held-out layers must never enter mechanistic claims; if a layer matters to the mechanism, measure it.

### 2.6 tanh saturation silences regulator columns (Table 1)

In the ground-truth universe the operating point can sit far from the tanh centre. The engine’s physical-state coordinates relate to the fitted model’s z-scored state (the ‘twin’) as x = μ + σ x_z, where μ and σ are the per-feature mean and standard deviation of the simulated trajectories in the generator’s own coordinates, computed NaN-aware over observed entries (§4.4). The ground-truth regulatory term tanh(μ_j + σ_j x_{z,j}) therefore carries the gradient factor σ_j·sech²(μ_j) with respect to x_{z,j} at the operating point (the steady input level at which the nonlinearity is evaluated, matched to the training distribution). Layer means μ can be extreme (observed maxima: G 46.6, P 278.1, S 26.8, M 25.5). At the strict numerical criterion |μ| ≥ 19 (sech²(μ) ∼ 10⁻¹⁶), the gradient backpropagated to the regulator state through each affected source column is approximately zero (P: 35; G: 7; S: 7; M: 2); the edge-weight gradients in that column need not vanish, because they contain the factor tanh(μ) ≈ ±1 rather than sech²(μ). The saturated input instead acts locally as a constant, making each affected weight contribution collinear with the corresponding target bias and obscuring their separate contributions. At the looser operating-point criterion |μ| > 3.0 (sech² < 0.01), the number of strongly attenuated columns rises to 127 in P (robust across thresholds 2.0–4.0, range 118– 139; G 62, S 84, M 20); all 127 fall within the 162 observed P features. Only features with defined μ are evaluable — fully unobserved features have undefined moments and are not scored as saturated. This is an operating-point-offset saturation, not a trajectory-wide lock-out (no feature saturates at 100% of observed time points). Numerically silent columns pose an identifiability obstacle, and strongly attenuated columns may limit achievable recovery. Saturation therefore constitutes the third measurable failure-mode component of the identifiability budget, alongside the oracle-init basin curve (§2.4) and corruption (§2.5); the ideal-budget ceiling test (§2.4) is a cross-cutting diagnostic rather than a budget component. We provide the saturation audit as a stand-alone tool (§4.9).

**Table 1.**
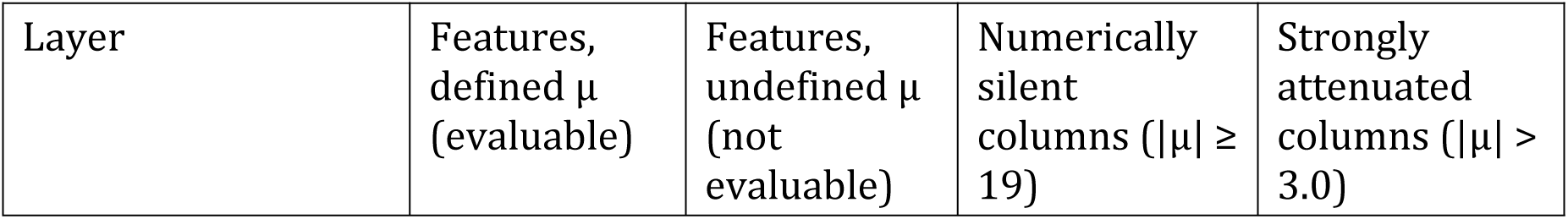
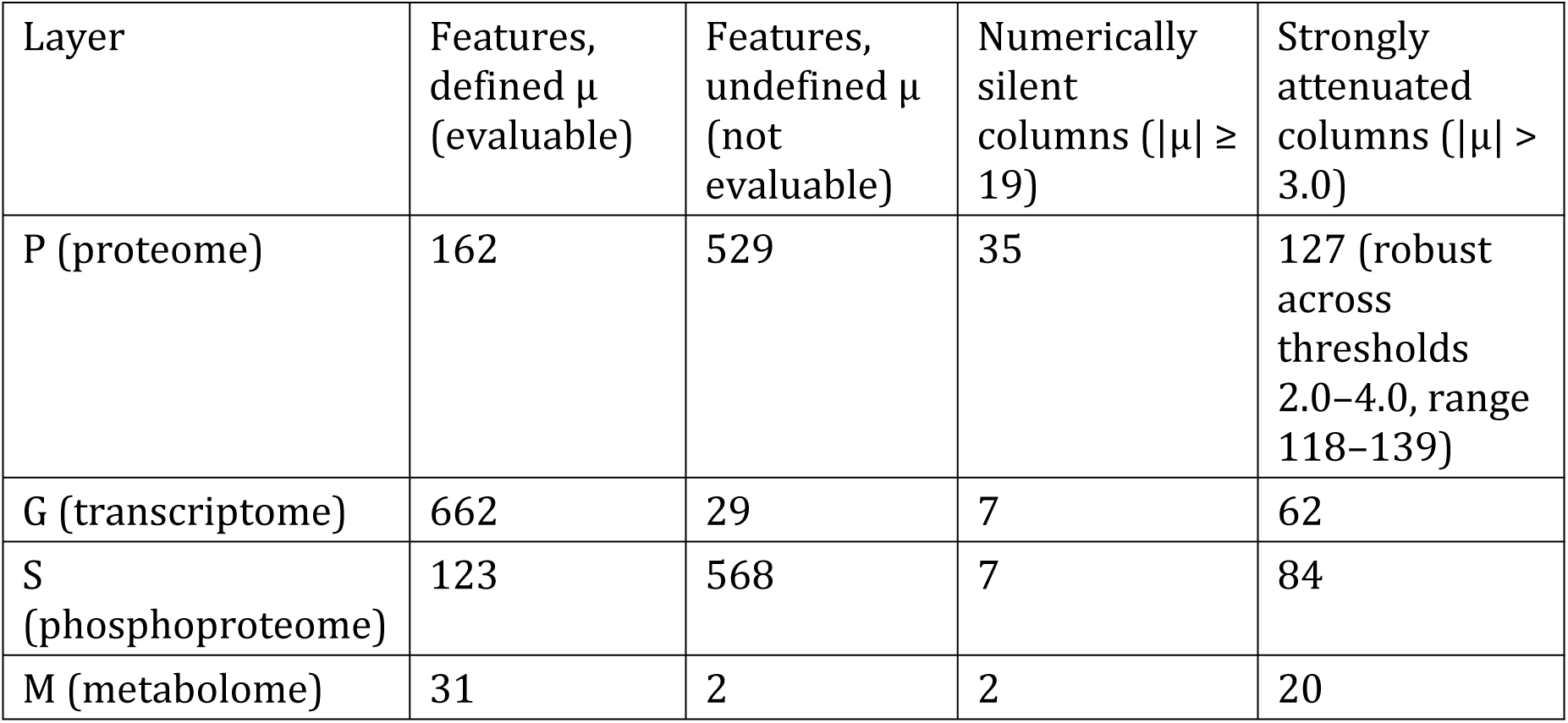
tanh-saturation column counts by layer and criterion (saturation audit, §2.6; numerically silent = regulator-state gradient numerically zero under the strict criterion, while edge-weight gradients need not vanish; strongly attenuated = sech² < 0.01 at the operating point, i.e. strongly attenuated but not numerically zero). Counts are evaluated within each layer’s features with defined μ; fully unobserved features have undefined μ and are not evaluable.

### 2.7 Operational semantics: the fitted twin executes interventional queries (Fig. 5)

Before trusting such queries on real data, we measured what they are worth in the ground-truth universe: we deleted the same two edges, both realized outgoing regulatory edges of Srebf1 (to Trp53 and Stat3), in the generator and in the σ = 0.1 oracle-init twin (§2.4). We compared the resulting knockout (KO)-minus-wild-type (WT) difference trajectories from a shared initial state. Intervention queries operate within the per-prediction confidence-grading framework (Q = one-hop, non-convergent; D = directional-only; N = not reportable; X = structurally unreachable; §4.6) — they interrogate a fitted twin, they do not amount to causal-edge identification.

**Figure 5.**
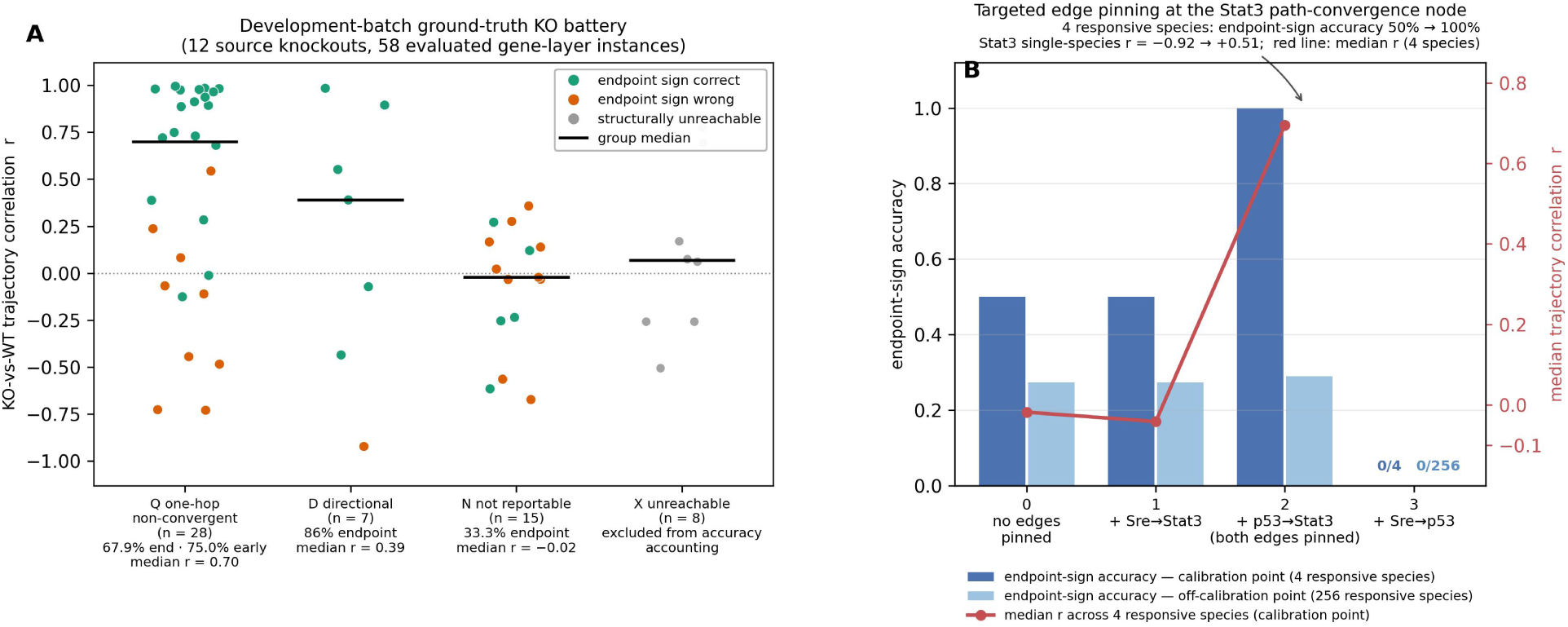
Intervention fidelity: knockout (KO) battery and targeted edge pinning. A: Development-batch ground-truth knockout (KO) battery (an independent-source batch of the same universe is defined in §4.6) — 58 evaluated gene-layer instances from 12 source knockouts (50 scored, 8 structurally unreachable), graded per §4.6 (Q n=28, D n=7, N n=15, X n=8); n denotes evaluated source–target instances; X is excluded from accuracy accounting. Points are per-instance KO-versus-wild-type (WT) trajectory correlations (colour: endpoint-sign correctness); horizontal bars are group medians. Q = one-hop, non-convergent; D = directional; N = not reportable; X = structurally unreachable. B: Targeted edge pinning at the Stat3 path-convergence node — bars: endpoint-sign accuracy at the calibration point (the operating point of §2.6; 4 responsive species, dark) and at one off-calibration point (256 responsive species, light); ‘responsive’ denotes a maximum ground-truth |KO−WT| difference across the trajectory exceeding 0.05 in z-space; red line: median r across the 4 responsive species at the calibration point (right axis). Manually pinning the node’s two incoming edges (Sre→Stat3, p53→Stat3) to their ground-truth weights restored it (Stat3 single-species r = −0.92 → +0.51; endpoint-sign accuracy 50% → 100%; the arrow marks the restored two-edge configuration). In the over-pinning control, pinning a third edge (Sre→p53) abolished the response (0/4 calibration, 0/256 off-calibration); r is undefined because the fitted difference trajectories became numerically constant. Calibration and off-calibration sets are each a single initial state; n denotes responsive species, not replicate experiments.

Three findings delimit the value of these queries. (i) The machinery is not the bottleneck: a ground-truth-parameter arm constructed with the first-order z-conjugate transform (no fitted-parameter error) matches the true intervention response in trajectory shape and endpoint sign at the calibrated operating point (defined in §2.6) (flattened r = 1.000, with trajectory matrices flattened to vectors before correlation; endpoint-sign agreement 100%). Even this ground-truth-parameter arm has a median max-abs amplitude ratio of 1.38 across the four responsive species (model versus ground truth); amplitude calibration is therefore not part of this shape-and-sign conclusion. (ii) The bottleneck is fitting error, and fidelity decays with network distance. A source–target instance denotes one knockout source evaluated against one target. Pooling 12 single-source knockouts (58 evaluated gene-layer instances; 50 scored and 8 structurally unreachable), one-hop targets on non-convergent paths recover trajectory shape and endpoint direction (n = 28; median trajectory r = 0.70; endpoint-sign accuracy 67.9%, early-window (§4.6) 75.0%), whereas two-hop effects are not reportable: development-batch trajectory correlations were near zero (median r = −0.02), the development-batch endpoint-sign accuracy was 33.3% against the 50% binary chance level (n = 15), and the independent batch contributed only six N-grade instances (5/6 endpoint-correct). The development and independent-source batches are two independently drawn sets of knockout source genes from the same ground-truth universe, not animal batches or experimental processing batches. Between these extremes, the 7 first-hop instances at convergent/cancellation nodes (grade D) show endpoint-direction agreement in 6/7 development-batch instances but poor trajectory-shape agreement (median r = 0.39); they are reported direction-only, with magnitude withheld (§4.6). Their trajectory-shape collapse is consistent with the competing-paths mechanism of finding (iii). (iii) Residual failures localize to path-convergence nodes whose net effect is a difference of competing paths: Stat3 integrates direct inhibition by Srebf1 with indirect disinhibition via Trp53, and a 26% weight error (relative, fitted versus ground truth on the matched-form σ = 0.1 oracle arm) on the direct edge reverses its predicted response (r = −0.92). As a diagnostic, we manually pinned this node’s two incoming edges to their ground-truth weights, and this alone restored it (exploratory, n = 4; endpoint-sign accuracy 50% → 100%; Stat3 trajectory r = +0.51 at the calibration point (the operating point of §2.6), and from −0.38 to +0.28 in the off-calibration arm (a second initial state, away from the calibrated operating point; 256 responsive species, i.e. those whose maximum ground-truth |KO−WT| difference across the trajectory exceeds 0.05 in z-space)).

This targeted correction is a direct quantification of what a single perturbation experiment buys. In the over-pinning control, pinning a third edge abolished the response (Fig. 5). Two intuitive confidence signals did not work: ensemble consensus across noise levels and robustness to ±20% weight perturbation both failed to separate correct from incorrect predictions. In the off-calibration condition, small-magnitude effects are anti-correlated even for the ground-truth-parameter arm. Calibrated confidence for deep-chain predictions thus remains open, a caution that parallels recent evidence that deep perturbation predictors can fail to outperform simple baselines when evaluation is not carefully instrumented [15].

A fitted white-box twin is an executable object: edges can be deleted and the system re-integrated to produce intervention-response trajectories. This capability distinguishes a dynamic twin from a static association model and provides the operational basis for the interventional reading of “causal” used throughout this paper.

### 2.8 Rank-level recovery holds across the Hill family; weight magnitudes do not (Table S2)

The remaining sections assemble two tiers of generality evidence: kinetic-family robustness (§2.8–2.9) and exploratory real-cohort re-analysis (§2.10). In the Hill-family arms, the ground-truth generator used Hill regulation (saturating Hill-equation kinetics) while the twin retained tanh kinetics — a cross-family mismatch test (candidate topology, edge set, observation budget and scoring protocol unchanged). Edge-ranking point estimates were above 0.5 across both oracle-initialized and cold-started Hill-family arms (tie-averaged (§4.6) pooled AUROC; Table S2), whereas recovered weight magnitudes agree only weakly with ground truth (signed Spearman 0.16–0.34; magnitude-only ≈ 0), consistent with amplitude distortion from L1 shrinkage toward zero during training. A descriptive account — sparse blocks move, dense blocks do not — reconciles all ten arms (Hill: s42, s43, s44 (dt0.25), basin, prior and blindv2; tanh: blind3, blind4, prior3 and prior4; basin reuses the seed-42 configuration and is not an independent replicate; Table S2) without arm-specific exceptions.

### 2.9 Delayed-family behaviour: integration-step stability and initialization sensitivity (Fig. S1, Fig. S3)

As a second kinetic-robustness test, the delayed-family arm asks whether edge recovery survives when regulation is transmitted with a fixed lag rather than instantaneously. It implements the §4.3 regulation through single-dimension relaxation (single coordinate) with a fixed lag (step conversion per §4.5); topology, candidate set and scoring are unchanged. Rank-level recovery of the oracle-initialized delayed-family GE arm holds under integration-step halving (AUROC 0.667/0.691/0.693 for dt 0.5/0.25/0.125; Fig. S3); the base arm is dt = 0.5, not a separate fourth condition. However, recovery is sensitive to initialization perturbation, with all noisy delayed-family initializations remaining near chance (Fig. S1) — consistent with the weak weight-level recovery reported above (§2.8). The two curves are not directly comparable: the oracle-init basin reports edge-class AUROC across the full ODE model, whereas the delayed-family init-sensitivity curve is a single GE-layer arm under a different kinetic family.

### 2.10 Exploratory differential analysis of a real five-omics cohort (rat endurance training)

As an exploratory real-cohort analysis outside the simulated benchmark, we re-analysed the MoTrPAC (Molecular Transducers of Physical Activity Consortium) rat endurance-training dataset [25] with a five-omics cross-sectional design (45 animals; cohort definition and estimand in Methods). Covariate-adjusted differential analysis (the primary group contrast of the 9 eight-week trained animals (completers) versus the remaining 36 animals — a mixed comparison group spanning all other training durations and the controls — §4.8, with sex as the sole covariate, run separately for each of the five omics layers) yielded significant hits (adj. p < 0.05): 33 transcripts, 1 protein, 0 phosphosites, 21 acetyl sites and 100 metabolites (numbers in Table S3). The metabolome shows the largest response spectrum. The sex main effect was markedly larger than the training effect at the transcript layer (8,361 vs 33 hits), consistent with sex as a dominant biological axis in this dataset. We report the covariate-adjusted contrast as primary. A sex-stratified listing and a nine-versus-nine sensitivity analysis restricted to eight-week trained versus eight-week control animals accompany the primary contrast in the Supplement (Supplementary Note S1, Tables N1a and N1b; stratified results listed for transparency, no hypothesis testing). Acetyl sites are an additional fifth-omics readout available in the rat dataset; the four-layer mouse framework has no corresponding layer.

## 3. Discussion

The intervention dissection of §2.5 admits a pressure-redistribution (“water-bed”) reading — the two bottlenecks are partially overlapping: pinning the bias channel recovers the larger share of the P-layer deficit (R² −8.8 → −5.2), whereas pinning the P-pathway raises R² to −6.6. Its weaker recovery relative to pinning bias alone is consistent with the over-constraint introduced by fixing translation coupling, bias and decay. Whether this compensation structure generalizes beyond the tested model class remains to be determined.

These results, together with the negative findings above, argue against treating trajectory fit as sufficient evidence for a mechanistic claim. For similar dynamic multi-omics models, we recommend reporting a static-baseline comparison together with the four audit instruments introduced in this study. Cold starts should be reported separately from oracle-initialized diagnostics so that an unattainable upper bound is not mistaken for a deployable result. Mechanistic claims should be restricted to measured layers and interpreted in light of these audits. Even an imperfectly recovered twin can still provide falsifiable hypotheses for targeted perturbation follow-up.

We define causal edges in the interventional sense (perturbing the source in the biological system changes the target’s dynamics) because trainable dynamic models answer what-if trajectory queries, not graphical conditional independencies. For the drug-mechanism questions studied here, the interventional notion is central because it concerns how perturbations alter the system. The benchmark question is therefore not “can a causal graph be discovered from omics data”, but “to what extent does fitting time-series dynamics identify intervenable causal edges across observation and optimization budgets?” The white-box design makes this definition operational: every learned edge is an explicit, deletable, re-testable hypothesis (§2.7). We therefore report intervention predictions with per-prediction confidence grades, within the interventional scope defined above.

The ground-truth intervention assay (§2.7) converts our budget thesis into an operational rule: in the σ = 0.1 oracle-init setting of §2.7, observational time courses provide preliminary one-hop trajectory-shape and direction information (within the Q-grade, with an explicit amplitude warning). Deep-chain fidelity is recovered edge by edge, and the edges with the highest payoff are the incoming edges of path-convergence nodes — precisely the nodes that dominate compensatory responses in drug mechanisms. A twin fitted on observational data is therefore not a finished predictive model. It is better read as a prioritized, payoff-quantified list of perturbation experiments; the twin’s interventional output and its calibration value are two aspects of the same analysis. This also disciplines claims about compensatory buffering: compensation is almost by definition a path-convergence phenomenon, and our assay shows its net sign is exactly what observational fitting cannot pin down (Stat3, §2.7). Compensation can be detected qualitatively, but quantifying it requires the targeted perturbations the twin itself nominates.

We position our work against two recent lines of work, developed independently and concurrently with ours. Our estimand is edge-level identifiability in multi-trajectory, trans-omics dynamics under insulin stimulation—the extent to which fitting dynamics identifies intervenable causal edges across observation and optimization budgets. This is complementary to cell-level virtual-perturbation models such as RegVelo [26], which nominate regulators that bias cell fate along single-cell trajectories. The estimands differ: ours audits edge-level mechanistic identifiability, whereas theirs predicts cell-level responses. Likewise, a fitted twin that executes in silico intervention queries is an instance of an individual-scale world model in the sense articulated by Xing and Song [27] and Noori et al. [16]. Our contribution is the measurement instrument that audits what such models recover, not another generative model. In our setting, the metabolite layer serves as a downstream readout of the multi-layer response; the mechanistic question is whether the intermediate edges linking an upstream perturbation to that readout are correctly identified, not merely whether the observed trajectories are fitted.

As noted in the Introduction, Giampiccolo et al. [22] provide a local, parameter-level account of identifiability. Our study instead examines edge-level structure recovery in a white-box masked ODE. We differ in three respects: (i) we treat identifiability as an operational, empirically measurable budget (the oracle-init basin curve, held-out-layer corruption localized by an intervention-based carrier decomposition, and tanh saturation ceilings), rather than a binary local property; (ii) controlled intervention experiments (zf, x0, bias and P-pathway pinning) on ground-truth networks turn the compensation between neural-network and mechanistic terms that they report qualitatively into a quantitatively attributed failure account with actionable prescriptions; and (iii) our semi-synthetic design extends the GeneNetWeaver/DREAM tradition [11,13] from single-layer gene-regulatory networks to four-layer trans-omics dynamics.

Single-cell perturbation-response models (scGen, GEARS, CPA and successors [28–30]) address a neighbouring but distinct problem: they predict population-level responses in learned embedding spaces, whereas our instruments audit edge-level identifiability against the exactly known ground-truth network of our semi-synthetic benchmark. The two are complementary: embedding-space response prediction can succeed while edge recovery fails, and the confidence triage of §4.6 characterizes the reliability of edge-level predictions when recovery is incomplete. The recent benchmark evidence that deep models trained on observational data do not yet outperform linear baselines in predicting unseen perturbation effects [15] underscores that this distinction is practical, not semantic.

Across the practically attainable budgets tested, fitting did not yield reliable edge recovery; substantial edge-ranking recovery was confined to unattainable oracle-initialized upper-bound diagnostics on measured layers. The binding constraint is therefore not trajectory fit alone but operational identifiability—an identifiability budget. What that budget does buy, in the same σ = 0.1 oracle-init setting, is preliminary one-hop trajectory-shape and direction information, with deeper structure purchased edge by edge through interventional follow-up; amplitude calibration is not established. These results support a limited practical use for the twin: prioritizing perturbation experiments for follow-up validation.

The multi-component regime is a relevant testbed for these methods: TCM formulas act on multiple targets and can trigger cascading responses across multiple omic layers. The present single-perturbation benchmark is the calibration experiment that supplies the missing measurement instruments. The TCM multi-component case is the next stress test for these instruments. A companion study applying them to multi-component intervention modeling is in progress.

### Limitations of the study

Our instruments were evaluated on a single trans-omics topology and a simplified model class with linear production, first-order decay and tanh regulation. The generator and the twin also share the same functional form, so the basin, held-out-corruption and saturation accounts are calibrated primarily to this matched-form setting. The Hill-family audit supports rank-level recovery under a functional-form mismatch, but does not establish these accounts for other functional forms or topologies.

The evaluation has five further boundaries. The metabolite layer contains only 33 species, so conclusions for that layer should be read with this coverage in mind. Baseline and optimization coverage remains limited: FD + LASSO and dynGENIE3 were tested, additional methods such as BINGO [31] remain untested, and the proximal regularization strength was not exhaustively scanned. We also did not test data-driven static pre-training as initialization, leaving open whether it could mitigate cold-start failure. The knockout battery is enriched for high-out-weight sources (56th–96th percentile among hub-eligible genes), so confidence calibration for weakly connected sources remains untested. Finally, edge recovery is evaluated only within the prior candidate set; de novo discovery beyond that set is outside the scope of this benchmark.

## 4. Methods

### 4.1 Data and preprocessing

We used the mouse-liver insulin-response trans-omics time series of Matsuzaki et al. [23] (transcriptome: GEO GSE166336; proteome and phosphoproteome: ProteomeXchange PXD022728 (proteome) and PXD022823 (phosphoproteome), PRIDE repository, LTQ Orbitrap Velos; metabolome: Tables S2–S3 of the source publication — to our knowledge no public repository accession exists for the metabolome layer). It profiles four molecular layers (transcriptome G, proteome P, phosphoproteome S, and metabolome M) under insulin and control (PBS) conditions. Each omics block retains its native sampling grid from the source publication (transcriptome and proteome: 0, 60, 120, 180, 240 min; phosphoproteome: 8 time points; metabolome: densely sampled at 0–30 min (0, 2.5, 5, 7.5, 10, 15, 20, 30 min) with additional measurements at 45, 60, 120, 180, and 240 min in the source study; the engine places metabolite trajectories on the full 13-point tissue-collection schedule (0–240 min)). No resampling to a common grid was performed. Every feature was z-scored in a NaN-aware manner (mean and standard deviation computed over observed values only). No imputation was applied to any observed data; only metabolite reference trajectories were linearly interpolated at unmeasured timepoints following the source publication’s pipeline, and interpolated points are excluded from loss and evaluation by the observation masks. Remaining missing values are likewise excluded through per-feature observation masks. Pathway-anchor mapping (mapping each feature to its KEGG pathway gene) was used for structural compression, defined here as intersecting the union of all KEGG KGML pathway nodes with the set of quantified gene products and detected metabolites to retain the measured features represented in that pathway union. Of the 14,076 quantified gene products (and 193 detected metabolites) before structural compression, 594 signaling genes form the candidate set. The working system comprises 2,106 molecular species (691 per molecular layer ×3 + 33 metabolites; the remaining 97 transcripts are modelled as targets but enter no candidate set).

### 4.2 Candidate prior masks

Candidate regulatory edges were assembled from prior knowledge only, as binary masks; no mask entry was derived from the time-series data themselves. Gene–gene regulatory edges (GE) combine KEGG KGML pathway relations with literature-curated transcription-factor targets (TRRUST v2 [32], mouse, each supported by at least one PubMed ID). Signalling-to-gene edges (SG) mirror the GE candidate topology, with the driving source replaced by the regulator’s phosphorylation activity; no new prior is introduced. Enzyme–metabolite edges (EC) combine KEGG KGML assignments with edges recovered by parsing reaction equations. Metabolite– metabolite edges (MM) require pathway co-occurrence in at least two KEGG pathways plus a small set of module-level priors. Feedback edges (FB) connect the five measured signaling metabolites retained in the simulated universe to the 594 signaling genes of the candidate set. The remaining measured metabolites were excluded because they fall outside the local KEGG KGML pathway union that defines the subnetwork. Protein–phosphosite edges (PS) form a dense candidate set within the 594 signaling genes of the candidate set, allowing every signaling protein to regulate the phosphosite state of every gene in that set. Within the sliced semi-synthetic candidate domain, the six masks contained 362 GE, 362 SG, 1,672 EC, 442 MM, 2,970 FB, and 352,836 PS edges (358,644 in total; PS = 594², including diagonal entries); class-wise retention probabilities are given in §4.4. In total the ground-truth network comprises 4,912 edges across six relation classes; every trainable weight is confined to its class mask, and all weights outside the masks are identically zero at all times. The two metabolite nodes with non-finite trajectories in the simulator (C00092, C00122) were excluded from metabolite-layer scoring.

### 4.3 White-box masked dynamics

Each modeled species obeys a first-order production–decay equation. Protein production is coupled linearly to transcript abundance through a softplus-calibrated per-gene translation term; regulation enters as bounded (tanh) summaries of masked upstream regulators (SG edges share the GE candidate mask by design, with the driving source replaced by phosphorylation activity); per-species additive constants (the bias channel) and first-order decay complete each equation, and an exogenous insulin pulse u(t) acts on the signalling layer during t ≤ 10 min (u(t) = 0 under PBS); u(t) is a coarse-grained signal-layer receptor pulse. The twin’s state x_z is z-scored; the ground-truth engine’s physical coordinates satisfy x = μ + σ x_z with per-feature raw-scale moments μ and σ (§2.6). In equations (G, P, S, M are the z-scored layer states; W the trainable weights; the class masks; d the per-layer decay constants; β the per-gene translation couplings (with its mask); b the bias channel):

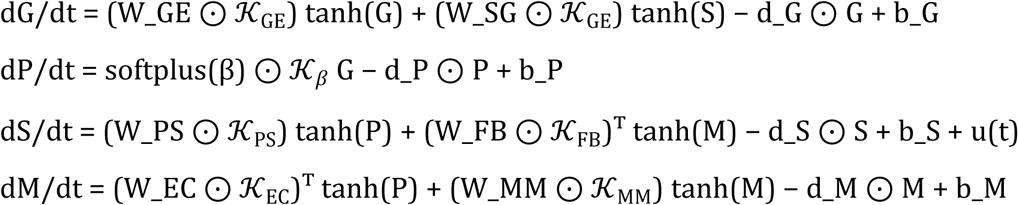

Trajectories are integrated with fixed-step RK4 (dt = 0.5 min, horizon 240 min) and training gradients are obtained by backpropagation through the solver (the adjoint method). The model is white-box: every parameter is an interpretable kinetic quantity (an edge weight, a decay constant, a translation coupling, an additive bias, or a signal-layer stimulus amplitude). The functional class — linear production gated by a softplus-calibrated layer coupling, tanh regulation, first-order decay — is the minimal skeleton shared by widely used trans-omics kinetic models (see §3, generality boundary).

### 4.4 Semi-synthetic ground-truth engine

To obtain a universe in which every causal quantity is exactly known, we instantiated the same model class as its own ground-truth engine. A ground-truth network was subsampled from the candidate masks (retention probabilities: GE, SG, MM, FB 0.3; EC 0.1; PS 0.01; fixed seed 42), yielding 4,912 realized edges with weights drawn from fixed magnitude–sign distributions. Reference ODE trajectories are deterministic under both conditions; three independent observation-noise realizations were then added and averaged per condition. Synthetic observations were generated by sampling the simulated trajectories on the empirical time grid, adding Gaussian observation noise (per-replicate σ = 0.1 in z-score space), and applying the observation mask (feature-level missingness matched to the real data: 76.6% proteome, 82.2% phosphoproteome). Under this missingness, 162 of the 691 proteome features carry at least one measurement under both conditions (insulin and PBS); ‘observed P features’ throughout refers to this set. An ideal-data mode (49-point 5-min grid, no missingness, no noise) provides the information-ceiling reference arm. Feature-wise z-statistics are stored separately and are regenerated during data preparation; fully unobserved features are stored as NaN and are excluded from all computations by the observation masks. Metabolite trajectories follow the full 13-point tissue-collection schedule; timepoints not measured in the source study are linearly interpolated, mirroring the source publication’s pipeline. All downstream claims about identifiability, fidelity, and confidence grading are made inside this universe and are labeled as such.

### 4.5 Training protocol

Unless stated otherwise, models were trained with Adam (learning rate 1e-3, 300 epochs). The seven loss terms are: masked trajectory MSE on each of the four layers (equal weight across layers), an L1 sparsity term (w_sparse = 1e-4), a direction-prior term penalizing sign-violating edges where the prior carries sign information (GE and EC classes only; w_dir = 1e-2), and a trajectory smoothness term on second differences (w_smooth = 1e-5). The sparsity weight was additionally scanned over two further regularization-weight settings (1e-3 and 1e-2, the w_sparse = 1e-3 and 1e-2 arms, both from zero initialization; Fig. 2A) to confirm that the failure mode under study is insensitive to the regularization weight.

Two sparsity regimes were compared. (i) Penalized only — the L1 term above with the proximal step disabled (w_prox = 0); on the real-data fit this retains all 3,878 candidate GE edges. (ii) Penalized plus proximal — after every optimizer update, a soft-thresholding operator w ← sign(w)·max(|w| − τ, 0) is applied to the six class-masked edge-weight matrices (GE, SG, EC, MM, FB, PS); translation couplings, decay constants, biases and the stimulus amplitude are not proxed, and in the frozen-block probe arms the frozen matrices are excluded from both gradient updates and the proximal step. Two proximal settings were run on the real-data fit (600 epochs each): τ = 3×10⁻⁴ together with w_sparse = 10⁻³, and τ = 10⁻³; across the two settings, 100–119 of the 3,878 candidate GE edges are retained.

Gradients were clipped to global norm 1.0; any step producing a non-finite gradient was skipped, and checkpoint saves were accepted only after a double finite-check of parameters and losses. Model selection uses the checkpoint from the epoch with the lowest training loss, which is saved during training and reloaded for all downstream evaluations; no validation-set selection is used because all arms are evaluated against ground truth, not predictive fit. For the frozen-edge probe arms (Fig. 4), the target weight matrices were frozen by setting their gradients to None before clipping, so the optimizer skips them and the proximal operator does not act on them. z-conjugate initialization uses the first-order conjugate transform of the dynamics (decays unchanged; layer couplings rescaled by the layer-wise standard deviation from z-scoring; edge weights rescaled by the corresponding ratios of these standard deviations and the local sech² factor of the tanh nonlinearity; constant terms absorbed into biases). Dense random or prior-signed initializations at amplitudes ≥ 0.3 fail by two distinct, amplitude-dependent pathologies (trajectory explosion at high amplitude; gradient death through tanh saturation at moderate amplitude) and require magnitude ≈ 0.05 with fan-in normalization to train at all. The cold-start arm family is defined in §2.4 (five statistically independent training runs; the zero-init arm counted once); the basin arm reuses the seed-42 run and is reported alongside; it is not counted as an independent replicate. Delayed coupling in the delayed-family arms was implemented as single-dimension relaxation (single coordinate) with a universal step conversion (lag × dt_ckpt ÷ dt_run, rounded); three-step data (dt0.5/dt0.25/dt0.125 arms) were checked with a cross-grid consistency tolerance of 0.05; the base arm is dt = 0.5 and is not a separate fourth condition.

### 4.6 Evaluation protocol and confidence grading

Edge recovery is scored by AUROC, AUPRC, and top-k precision against the ground-truth network within the identical candidate set. All pooled AUROC values are computed with tie-averaged ranks: in near-chance arms, where large tied score blocks make the pooled statistic sensitive to within-tie ordering, naive-rank and tie-averaged implementations differ by up to 0.05. We report tie-averaged values throughout (naive-rank values are not shown). We adopted the following negative-control criterion for the delayed-family dynGENIE3 phase-scrambled control (§4.7): a scrambled-arm pooled AUROC < 0.58 and a real-minus-scrambled margin ≥ 0.02 were required to declare separable signal. For that control, rank-invariance diagnostics (checks that pooled AUROC is stable across score-implementation orderings) carry a negative-control exemption: arms that do not meet this separable-signal criterion are excluded from cross-implementation rank comparisons. Held-out-layer fit is scored by R² against a flat observation-mean baseline, reported separately per condition.

Because the edge classes are severely imbalanced (362 GE vs. 2,970 FB candidates), pooling edges across classes would produce a Simpson-type inflated AUROC; we therefore score edges within each edge-class candidate domain, a quantity we denote edge-class AUROC throughout. For edge-class AUROC, a reported range a–b denotes the minimum and maximum of the six per-class values; the primary benchmark does not pool AUROC across edge classes. Separately labelled pooled AUROC in the Hill-family (§2.8) and dynGENIE3 (§4.7) audits is a descriptive summary and is not directly comparable with edge-class AUROC. Intervention fidelity was assessed with a battery of 12 knockout source genes selected by a fixed rule. The rule: genes with ≥2 outgoing ground-truth GE edges, ranked by total absolute outgoing weight, top 12; sources span the 56th–96th percentile of out-weight among hub-eligible genes, a moderate hub bias that we report as a limitation. Each knockout yields per-target trajectory predictions scored by correlation, endpoint sign, and early-response sign (early window = the first 30 min of the trajectory) against the known truth; the 12 source genes contribute different numbers of evaluated targets, yielding 58 evaluated gene-layer source–target instances in total (50 scored, 8 structurally unreachable).

Using this battery, we calibrated a per-prediction confidence triage with four grades (Supplementary Fig. S2): **Q (one-hop, non-convergent)**: first-hop targets on non-convergent paths: trajectory shape and direction reported, with an explicit amplitude warning — effect amplitudes are systematically over-estimated (median max-abs amplitude ratio 5.84 in the development batch and 4.78 in the independent source batch; log–log calibration slopes 0.163 and 0.176) (endpoint accuracy 67.9%, early 75.0%, median trajectory r = 0.70, n = 28). **D (directional)**: first-hop targets at convergent/cancellation nodes: endpoint direction reported with an explicit warning, magnitude withheld (6/7 development-batch and 8/8 independent-source endpoint accuracy; development-batch Wilson 95% CI 49–97%; median r = 0.39, i.e. direction right, dynamics wrong; n = 7 development; D-grade max-abs amplitude ratios 9.31 (development) and 5.68 (independent), reported separately from Q). A sign-level follow-up test on the D-graded instances that have a realized direct edge (12 of 15 across the two batches) discriminates two competing explanations. A direct-only effect would carry the direct-edge sign; indirect paths may reverse the net endpoint sign. In all 12 cases the ground-truth endpoint is dominated by the indirect paths, and the fitted model reproduces these indirect-dominated endpoints in 11 of 12; the single exception is a development-batch instance whose fitted direct-only contribution (+17.82), indirect-only contribution (−7.06) and net (+10.75) carry the opposite sign to the ground-truth endpoint (−2.17). The remaining three D-graded instances have no realized direct edge and cannot be decomposed this way; the fitted model assigns each a non-zero direct edge. The D grade’s directional reliability therefore reflects genuinely captured net effects, while its trajectory collapse reflects dynamics error along the path, not sign confusion. **N (not reportable)**: targets two or more hops away. In the development batch, endpoint-sign accuracy was a below-chance point estimate without statistical separation from chance (n = 15; 10/15 near-zero endpoints, |true endpoint| < 0.001). The predominance of near-zero endpoints in the development batch limits the interpretability of its exact sign scores. **X (structurally unreachable)**: targets with no ground-truth path from the perturbed source, excluded from accuracy accounting (n = 8). Endpoint sign is scored by exact three-valued sign match (−1/0/+1): negative values map to −1, exactly zero to 0, and positive values to +1, with no rounding or thresholding in the sign assignment; near-zero but nonzero endpoints therefore retain their numerical sign. Scored endpoints are continuous, and no zero-sign instance occurs in the battery, so the effective chance level is 50%. Reported accuracies carry Wilson score 95% confidence intervals: one-hop endpoint-sign 67.9% (19/28, CI 49–82%), two-hop sign 33.3% (5/15, CI 15–58%), targeted edge pinning 100% (4/4, CI 51–100%), and one-hop early-window 75.0% (21/28, CI 57–87%). The Q/N boundary shows a borderline trend (endpoint accuracy 67.9% vs 33.3%; Fisher exact p ≈ 0.05, exploratory and unadjusted). D and Q do not differ in endpoint accuracy in the development batch (p = 0.64), and the D grade rests on trajectory-fidelity collapse.

Independent-source validation. The grading was further tested on an independent batch of knockout sources drawn from the same ground-truth universe (33 evaluated instances: Q 17 of 17, D 8 of 8, N 5 of 6, X 2; scored = 17 + 8 + 6 (N’s denominator) = 31). Endpoint accuracy differed significantly between batches for Q (Fisher p = 0.0086) and showed no significant batch difference for D (p = 0.4667, which does not establish equivalence); N carried no reliable direction evidence in the development batch; the independent batch’s six N-grade instances (5/6 endpoint-correct) are too few to support a stable accuracy estimate. Cross-universe validation remains open.

Grade thresholds are a first-pass empirical calibration on this single universe and will be updated as validation instances accrue; when hop distance is unknown (real universes), grades automatically degrade one step (Q→D). Per-prediction confidence, not per-model confidence, is the intended output unit. The below-chance reading is a development-batch point-estimate claim, supported by the convergence-node mechanism (finding (iii)). A species was scored as responsive when its maximum ground-truth |KO−WT| difference across the trajectory exceeded 0.05 in z-space; the calibration point carried 4 responsive species and the off-calibration point carried 256 responsive species (insulin condition). Sign scoring is implemented in the analysis pipeline (released as part of the evaluation code).

### 4.7 Static baseline

As a static baseline without dynamic priors (the candidate domain is delimited by prior masks) we implemented finite-difference derivative estimates followed by per-target LASSO regression (hand-written iterative shrinkage-thresholding, ISTA; λ selected by BIC) over the cumulatively sampled design matrix. AUROC is computed by a hand-written Mann–Whitney average-rank procedure. As an independent method control we additionally benchmarked dynGENIE3 [33], the dynamical time-series extension of the tree-ensemble method GENIE3 [34], on the GE candidate subset defined by the subsetting rule requiring observation under both conditions (228 of 239 GE-layer genes). We scanned two observation modes over the same ground-truth universe (2 modes × 3 tree settings × 5 seeds = 30 runs), scored by tie-averaged AUROC within the GE mask domain. The raw dynGENIE3 output is VIM[regulator, target]; scoring used its transpose, VIM^T[target, regulator], with both index directions checked. All cross-method comparisons in this paper are made within their respective candidate domains. The run plan specified 45 runs across three planned observation datasets. The third dataset was excluded because its observation files were identical to those of the first, leaving two observation modes over the same ground-truth universe and 30 runs for analysis. Run-level scores are reported in Table S4: dynGENIE3 attains 0.607 ± 0.016 (95% t-interval half-width, n = 5 seeds, Table S4) on sim2i at the 1000-tree setting (the ideal-data mode of §4.4) and 0.562 (mean across five seeds) on sim2 (the realistic observation mode over the same ground-truth universe, with the observation mask matched to the real data). At the scale of all candidate edges across the 2,106-node universe scored per edge, tree-ensemble methods remain computationally impractical and do not exploit trajectory differential structure. A phase-scrambled negative control on the delayed-family sequences showed no separable signal (tier means across three seeds: real AUROC 0.4985/0.4894 versus scrambled 0.5356/0.5206 at the 500-and 1000-tree tiers; Δ = −0.0371/−0.0312), indicating that dynGENIE3 variable-importance scores are uninformative on these short delayed sequences; delayed-family rank recovery is therefore reported from the twin’s own estimator alone. The phase-scrambled control was restricted to the delayed-family sequences; no corresponding scrambled control was run for the sim2i ideal-data mode.

### 4.8 Exploratory rat cohort re-analysis (MoTrPAC)

As an exploratory analysis, we re-analysed the public MoTrPAC rat endurance-training dataset [25] (five-omics cross-sectional design; liver; endurance-training cohort dataset motrpac_rat-training-06_liver_v1 (PASS1B release), MoTrPAC Data Hub, motrpac-data.org; accessed August 2026). Of the 147 animals registered in the study (123 exercise, 24 control), per-layer sample counts were 52 (transcriptome), 60 (proteome), 60 (phosphoproteome), 54 (acetylome) and 50 (metabolome). Their intersection, after five design exclusions (two of ten 1-week animals, one 4-week animal, one 8-week trained animal and one 8-week control; 10% of the eligible set, skewed toward early-timepoint exercise), comprised the 45-animal analysis set (36 trained (8/10/9/9 at 1/2/4/8 weeks) and 9 eight-week controls; 25 females, 20 males). For the transcriptome layer, 54 vials comprised 50 animal samples and 4 QC vials; deduplication yielded 52 unique 5-digit IDs: 50 animal IDs and 2 non-animal quality-control prefixes. The cohort was defined before analysis (50 animals → 45 by design exclusions; cohort flow in Supplementary Fig. S4). The primary contrast compares the 9 eight-week trained animals (completers) with the remaining 36 animals (the 27 animals trained for other durations and the 9 eight-week controls); a nine-versus-nine contrast (8-week trained vs 8-week controls) is reported as a sensitivity analysis (Supplementary Note S1). Differential analysis used a linear model per layer with the design matrix [intercept, group, sex] — sex the sole covariate, group the main effect — with Benjamini–Hochberg adjustment within each layer.

### 4.9 Availability

The complete code pipeline will be made publicly available on GitHub (https://github.com/zhulv-zhang/metabolic-twin-benchmark) and archived on Zenodo upon publication. To avoid redistributing KEGG content, raw KEGG downloads and KEGG-derived candidate-edge masks and data are not included; the pipeline provides scripts for users to obtain the required inputs and regenerate these artifacts locally. The final analysis-input files used in this study were verified to be unchanged between two independent builds (30–31 August 2026 and 13 November 2026); later KEGG versions may yield different derived files. Trained parameters and per-arm training logs are not included; commands and seeds used in the workflows are documented in docs/ARM_LIST.md. The subsetting rule that reduced the full index(14,076 gene products) to the working system is documented in §4.1. Data used in the preparation of this article were obtained from the Molecular Transducers of Physical Activity Consortium (MoTrPAC) database, which is available for public access at motrpac-data.org. Specific datasets used are motrpac_rat-training-06_liver_v1 (PASS1B release).

## Author contributions

Z.Z. conceived and designed the study, developed the benchmark and the four instruments, implemented the pipeline, performed all analyses, and wrote the manuscript. L.J. contributed to prior curation and data preparation. Y.P. supervised the study, acquired resources, and revised the manuscript. H.Z. acquired funding. Y.P. and H.Z. are the corresponding authors.

## Funding

Scientific and Technological Innovation Project of China Academy of Chinese Medical Sciences (CI2026A04501).

## Notes

### Competing Interest Statement

The authors have declared no competing interest.

### Summary of Updates

This revision updates the title, abstract, main text, figures, and supplementary materials; clarifies candidate domains and evaluation metrics; expands the benchmark and robustness audits; and updates the confidence-grading and availability statements.

## References

1. Yugi K, Kubota H, Hatano A, Kuroda S. Trans-omics: how to reconstruct biochemical networks across multiple “omic” layers. Trends Biotechnol. 2016;34(4):276–290. doi:10.1016/j.tibtech.2015.12.013. PMID:26806111.

2. Kawata K, Hatano A, Yugi K, Kubota H, Sano T, Fujii M, Tomizawa Y, Kokaji T, Tanaka KY, Uda S, Suzuki Y, Matsumoto M, Nakayama KI, Saitoh K, Kato K, Ueno A, Ohishi M, Hirayama A, Soga T, Kuroda S. Trans-omic analysis reveals selective responses to induced and basal insulin across signaling, transcriptional, and metabolic networks. iScience. 2018;7:212–229. doi:10.1016/j.isci.2018.07.022. PMID:30267682.

3. Kokaji T, Hatano A, Ito Y, Yugi K, Eto M, Morita K, Ohno S, Fujii M, Hironaka K, Egami R, Terakawa A, Tsuchiya T, Ozaki H, Inoue H, Uda S, Kubota H, Suzuki Y, Ikeda K, Arita M, Matsumoto M, Nakayama KI, Hirayama A, Soga T, Kuroda S. Trans-omic analysis reveals allosteric and gene regulation axes for altered hepatic glucose-responsive metabolism in obesity. Sci Signal. 2020;13(660):eaaz1236. doi:10.1126/scisignal.aaz1236. PMID:33262292.

4. Zhang K, Zhou HY, Baptista-Hon DT, Gao Y, Liu X, Oermann E, Xu S, Jin S, Zhang J, Sun Z, Yin Y, Razmi RM, Loupy A, Beck S, Qu J, Wu J; International Consortium of Digital Twins in Medicine. Concepts and applications of digital twins in healthcare and medicine. Patterns (N Y). 2024;5(8):101028. doi:10.1016/j.patter.2024.101028. PMID:39233690.

5. Alsalloum GA, Al Sawaftah NM, Percival KM, Husseini GA. Digital twins of biological systems: a narrative review. IEEE Open J Eng Med Biol. 2024;5:670–677. doi:10.1109/OJEMB.2024.3426916. PMID:39184962.

6. Zhang P, Zhang D, Zhou W, Wang L, Wang B, Zhang T, Li S. Network pharmacology: towards the artificial intelligence-based precision traditional Chinese medicine. Brief Bioinform. 2024;25(1):bbad518. doi:10.1093/bib/bbad518. PMID:38197310.

7. Cui G, Li M, Guo W, Gao M, Zhu Q, Liao J. AI driven network pharmacology: multi-scale mechanisms of traditional Chinese medicine from molecular to patient analysis. Comput Struct Biotechnol J. 2025;27:5087–5104. doi:10.1016/j.csbj.2025.11.016. PMID:41322006.

8. Rackauckas C, Ma Y, Martensen J, Warner C, Zubov K, Supekar R, Skinner D, Ramadhan A. Universal differential equations for scientific machine learning. arXiv:2001.04385. 2020. doi:10.48550/arXiv.2001.04385.

9. Karpatne A, Jia X, Kumar V. Knowledge-guided machine learning: current trends and future prospects. arXiv:2403.15989. 2024. doi:10.48550/arXiv.2403.15989.

10. Brunton SL, Proctor JL, Kutz JN. Discovering governing equations from data by sparse identification of nonlinear dynamical systems. Proc Natl Acad Sci USA. 2016;113(15):3932–3937. doi:10.1073/pnas.1517384113. PMID:27035946.

11. Marbach D, Costello JC, Küffner R, Vega NM, Prill RJ, Camacho DM, et al. Wisdom of crowds for robust gene network inference. Nat Methods. 2012;9(8):796–804. doi:10.1038/nmeth.2016. PMID:22796662.

12. Prill RJ, Marbach D, Saez-Rodriguez J, Sorger PK, Alexopoulos LG, Xue X, Clarke ND, Altan-Bonnet G, Stolovitzky G. Towards a rigorous assessment of systems biology models: the DREAM3 challenges. PLoS One. 2010;5(2):e9202. doi:10.1371/journal.pone.0009202. PMID:20186320.

13. Schaffter T, Marbach D, Floreano D. GeneNetWeaver: in silico benchmark generation and performance profiling of network inference methods. Bioinformatics. 2011;27(16):2263–2270. doi:10.1093/bioinformatics/btr373. PMID:21697125.

14. Hill SM, Heiser LM, Cokelaer T, Unger M, Nesser NK, Carlin DE, et al. Inferring causal molecular networks: empirical assessment through a community-based effort. Nat Methods. 2016;13(4):310–318. doi:10.1038/nmeth.3773. PMID:26901648.

15. Ahlmann-Eltze C, Huber W, Anders S. Deep-learning-based gene perturbation effect prediction does not yet outperform simple linear baselines. Nat Methods. 2025;22(8):1657–1661. doi:10.1038/s41592-025-02772-6. PMID:40759747.

16. Noori A, Fishman N, Fang A, Fesser L, Zitnik M. World models for biomedicine. Cell. 2026;189(19):5845–5869. doi:10.1016/j.cell.2026.08.032.

17. Dupire L, Khan AA, Karaletsos T, Kelley S, Lundberg E, Ma J, Paull E, Quake SR, Rabadan R, Rowan C, Sims P, Tavazoie S, Tsang JS, Zhang M, Califano A. Fifteen challenges for generative AI applications to cell biology. Cell. 2026;189(19):5870–5892. doi:10.1016/j.cell.2026.07.004. PMID:42607668.

18. Miao H, Xia X, Perelson AS, Wu H. On identifiability of nonlinear ODE models and applications in viral dynamics. SIAM Rev. 2011;53(1):3–39. doi:10.1137/090757009.

19. Raue A, Kreutz C, Maiwald T, Bachmann J, Schilling M, Klingmüller U, Timmer J. Structural and practical identifiability analysis of partially observed dynamical models by exploiting the profile likelihood. Bioinformatics. 2009;25(15):1923–1929. doi:10.1093/bioinformatics/btp358. PMID:19505944.

20. Villaverde AF. Observability and structural identifiability of nonlinear biological systems. Complexity. 2019;2019:8497093. doi:10.1155/2019/8497093.

21. Browning AP, Warne DJ, Burrage K, Baker RE, Simpson MJ. Identifiability analysis for stochastic differential equation models in systems biology. J R Soc Interface. 2020;17(173):20200652. doi:10.1098/rsif.2020.0652. PMID:33323054.

22. Giampiccolo S, Reali F, Fochesato A, Iacca G, Marchetti L. Robust parameter estimation and identifiability analysis with hybrid neural ordinary differential equations in computational biology. NPJ Syst Biol Appl. 2024;10:139. doi:10.1038/s41540-024-00460-3. PMID:39609454.

23. Matsuzaki F, Uda S, Yamauchi Y, Matsumoto M, Soga T, Maehara K, Ohkawa Y, Nakayama KI, Kuroda S, Kubota H. An extensive and dynamic trans-omic network illustrating prominent regulatory mechanisms in response to insulin in the liver. Cell Rep. 2021;36(8):109569. doi:10.1016/j.celrep.2021.109569. PMID:34433063.

24. Meinshausen N, Bühlmann P. High-dimensional graphs and variable selection with the Lasso. Ann. Statist. 2006;34(3):1436–1462. doi:10.1214/009053606000000281.

25. MoTrPAC Study Group; Lead Analysts; MoTrPAC Study Group. Temporal dynamics of the multi-omic response to endurance exercise training. Nature. 2024;629(8010):174–183. doi:10.1038/s41586-023-06877-w. PMID:38693412.

26. Wang W, Hu Z, Weiler P, Mayes S, Lange M, Fountain DM, Haug JO, Wang J, Xue Z, Sauka-Spengler T, Theis FJ. RegVelo: gene-regulatory-informed dynamics of single cells. Cell. 2026;189(12):3773–3800.e44. doi:10.1016/j.cell.2026.04.022. PMID:42119563.

27. Xing EP, Song L. A world model of the virtual cell. Cell. 2026;189(19):5831– 5844. doi:10.1016/j.cell.2026.08.042.

28. Lotfollahi M, Wolf FA, Theis FJ. scGen predicts single-cell perturbation responses. Nat Methods. 2019;16(8):715–721. doi:10.1038/s41592-019-0494-8. PMID:31363220.

29. Roohani Y, Huang K, Leskovec J. Predicting transcriptional outcomes of novel multigene perturbations with GEARS. Nat Biotechnol. 2024;42(6):927–935. doi:10.1038/s41587-023-01905-6. PMID:37592036.

30. Lotfollahi M, Klimovskaia Susmelj A, De Donno C, Hetzel L, Ji Y, Ibarra IL, et al. Predicting cellular responses to complex perturbations in high-throughput screens. Mol Syst Biol. 2023;19(6):e11517. doi:10.15252/msb.202211517. PMID:37154091.

31. Aalto A, Viitasaari L, Ilmonen P, Mombaerts L, Gonçalves J. Gene regulatory network inference from sparsely sampled noisy data. Nat Commun. 2020;11:3493. doi:10.1038/s41467-020-17217-1. PMID:32661225.

32. Han H, Cho JW, Lee S, Yun A, Kim H, Bae D, Yang S, Kim CY, Lee M, Kim E, et al. TRRUST v2: an expanded reference database of human and mouse transcriptional regulatory interactions. Nucleic Acids Res. 2018;46(D1):D380–D386. doi:10.1093/nar/gkx1013. PMID:29087512.

33. Huynh-Thu VA, Geurts P. dynGENIE3: dynamical GENIE3 for the inference of gene networks from time series expression data. Sci Rep. 2018;8:3384. doi:10.1038/s41598-018-21715-0. PMID:29467401.

34. Huynh-Thu VA, Irrthum A, Wehenkel L, Geurts P. Inferring regulatory networks from expression data using tree-based methods. PLoS One. 2010;5(9):e12776. doi:10.1371/journal.pone.0012776. PMID:20927193.

