## Supplementary materials for "Trajectory fit alone does not identify causal edges: a white-box masked ODE benchmark for trans-omics digital twins of drug action mechanisms"

### Supplementary Figures

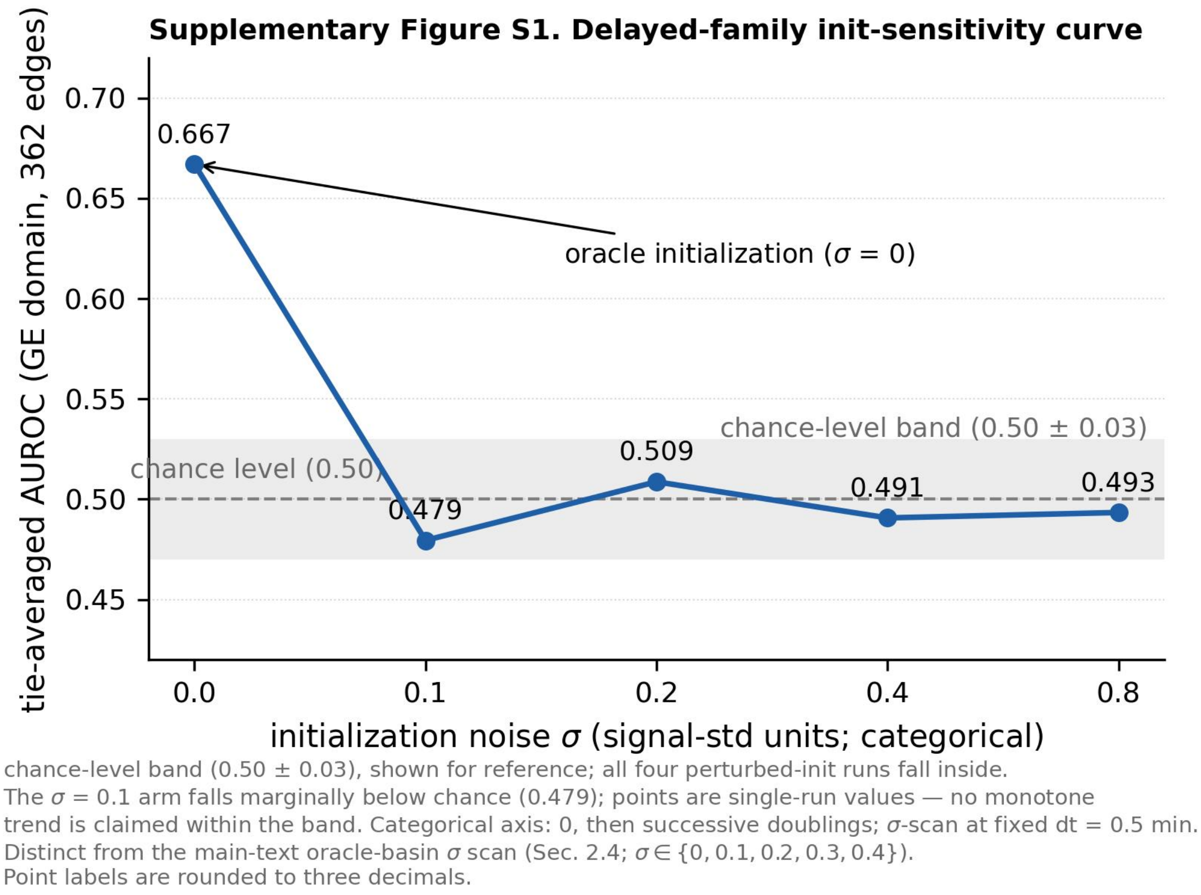


**Fig. S1.** *Delayed-family initialization sensitivity. Tie-averaged AUROC of the delayed-family GE arm within the 362-edge GE candidate domain under initialization perturbation. The main-text oracle-init basin (Fig. 3) uses a different kinetic family and scoring scope and is not directly comparable. Points are single-run values; point labels are rounded to three decimals. The shaded band marks chance level (0.50 ± 0.03), shown for reference; all four perturbed-init runs fall inside. Categorical axis: 0, then successive doublings; σ-scan at fixed dt = 0.5 min.*

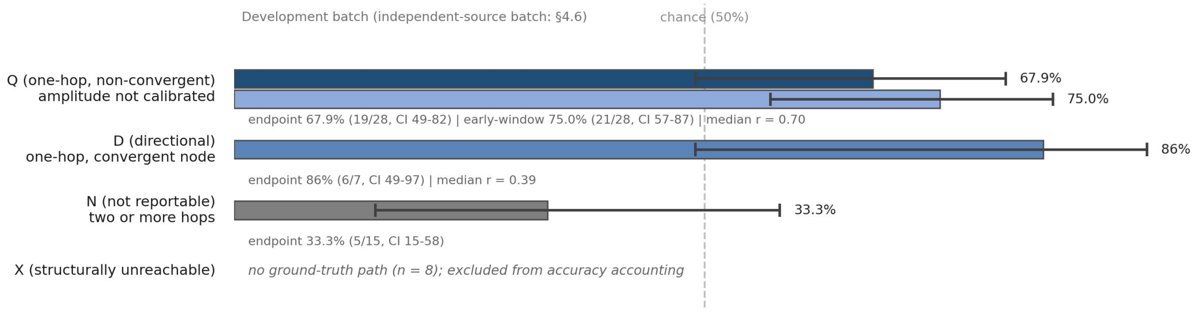


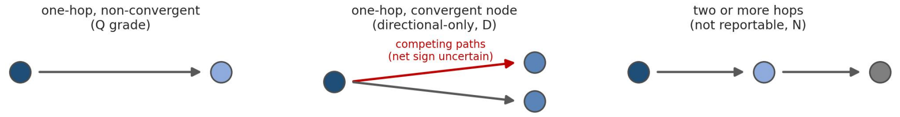


**Fig. S2.** *Per-prediction confidence triage (development batch). Q (one-hop, non-convergent): endpoint 67.9% (19/28, Wilson 95% CI 49–82%), early-window 75.0% (21/28, Wilson 95% CI 57–87%), median r = 0.70; amplitude is not calibrated. D (directional): endpoint 86% (6/7, Wilson 95% CI 49–97%), median r = 0.39; magnitude withheld. N (not reportable): endpoint 33.3% (5/15, Wilson 95% CI 15–58%). X (structurally unreachable): no ground-truth path (n = 8), excluded from accuracy accounting. The independent-source batch is reported in §4.6.*

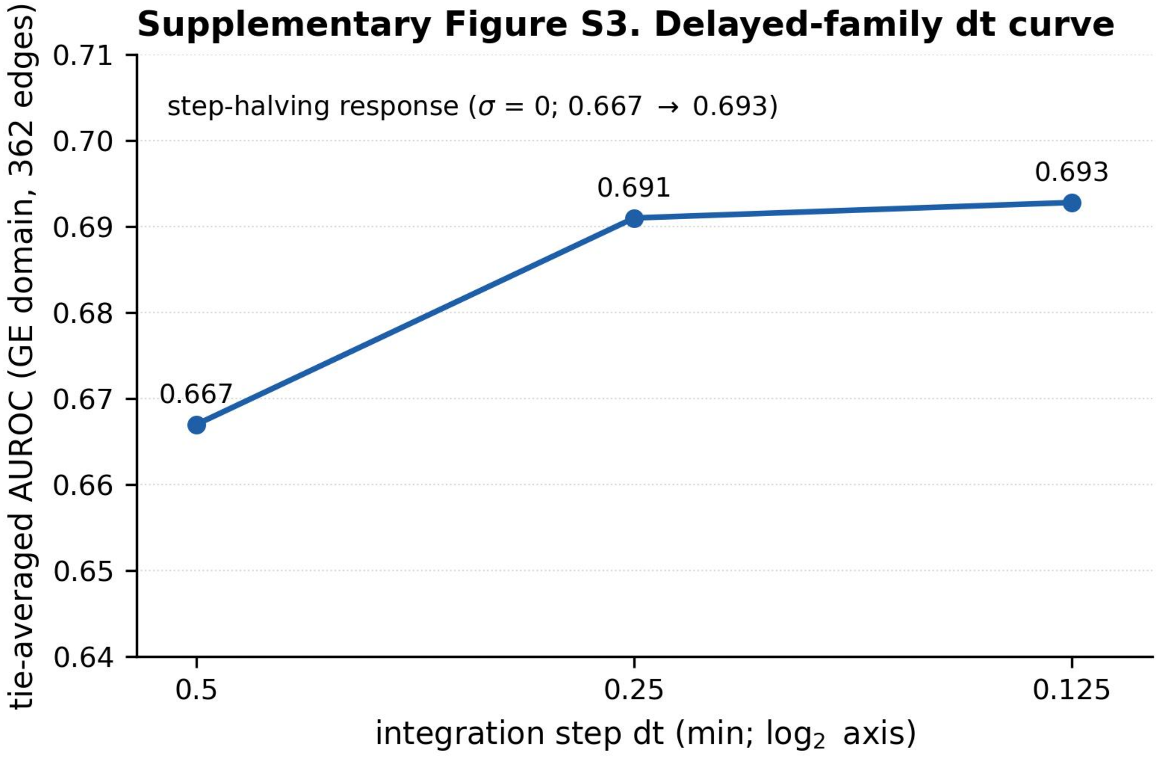


dt-scan at fixed $\sigma$ = 0 (oracle initialization); points are single-run values; tie-averaged convention per Methods. Point labels are rounded to three decimals.

**Fig. S3.** *Delayed-family integration-step stability. Tie-averaged AUROC of the oracle-initialized delayed-family GE arm within the 362-edge GE candidate domain under integration-step halving (0.667/0.691/0.693 for dt = 0.5/0.25/0.125 min). Points are single-run values; point labels are rounded to three decimals. The base arm is dt = 0.5 min and is not a separate fourth condition.*

### Supplementary Figure S4. Rat cohort flow

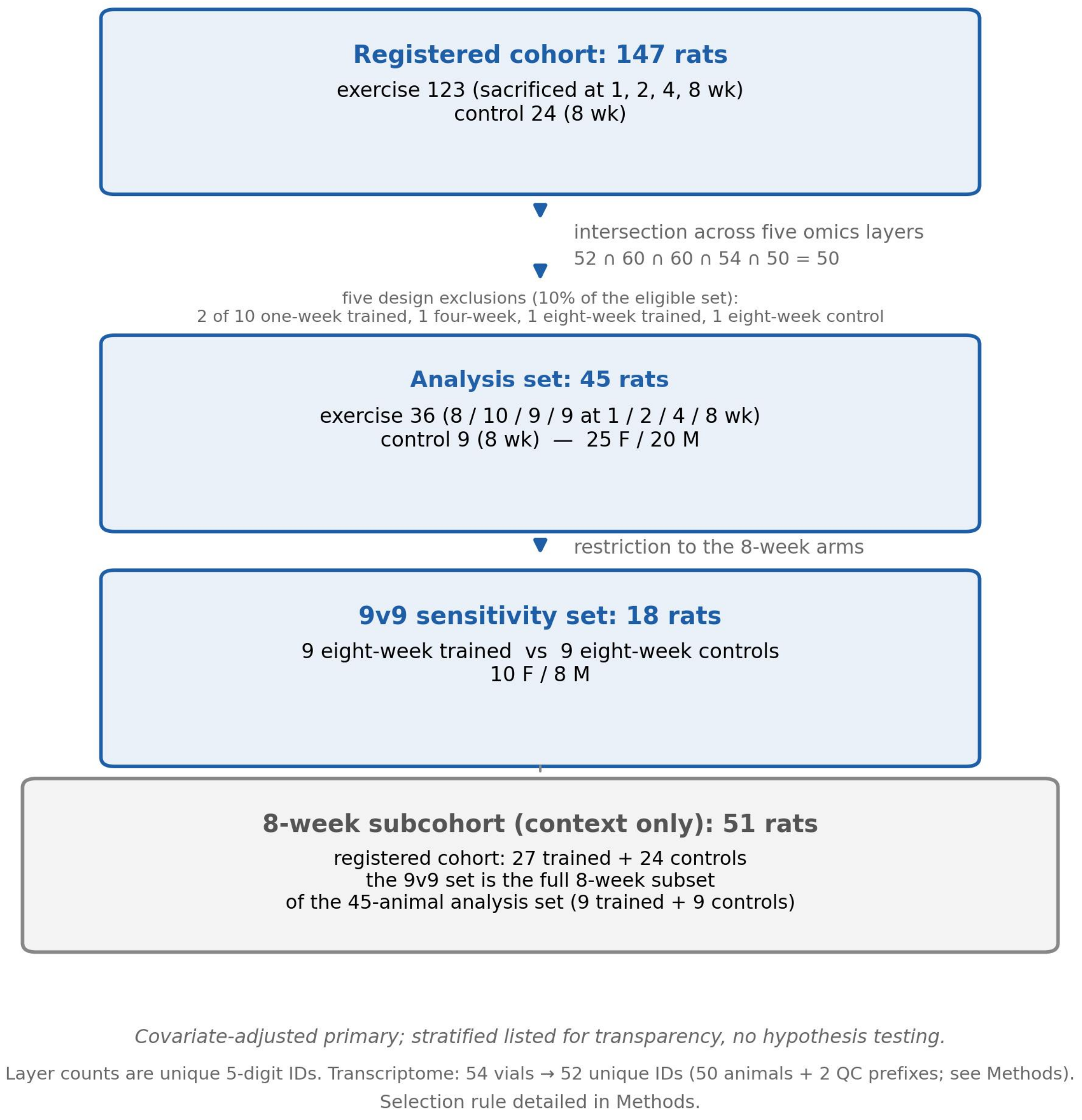


**Fig. S4.** *Rat cohort flow. Registered cohort (147 rats) → five-omics intersection (50 rats) → five design exclusions (10% of the eligible set) → analysis set (45 rats; 36 trained, 9 eight-week controls, 25 F/20 M) → 9-vs-9 sensitivity set (18 rats; 9 eight-week trained vs 9 eight-week controls, 10 F/8 M). The 8-week registered subcohort (51 rats) is background context only; the 9-vs-9 set is the full 8-week subset of the 45-animal analysis set.*

### Supplementary Tables

#### Table S1. Four completed intervention arms of the zf-arm dissection (held-out proteome and phospho-signalling layer $R^2$).

| Arm | Intervention | P-layer $R^2$ (two-condition mean) | Insulin | PBS | Note |
|---|---|---|---|---|---|
| zf (reference) | none (freeze-probe) | −8.8 | −8.89 | −8.73 | S layer −0.13 (−0.09/−0.17); held-out layers P and S; observed layers fit in-sample (G 0.39; M 0.85–0.86) |
| x0 | freezing the learnable initial state | −10.8 | — | — | makes the P layer worse; excluded as a remedy |
| bias | pinning the bias channel | −5.2 | −5.19 | −5.13 | largest recovery |
| P-pathway | pinning the P-pathway coefficients | −6.6 | −6.62 | −6.51 | second-largest recovery |

**Note. $R^2$ values are two-condition means with per-condition values in the Insulin and PBS columns where available.**

#### Supplementary Table S2 · Ten-arm pooled AUROC (kinetic-family audit)

| Arm | Family | Pooled tie-averaged AUROC | GE-domain AUROC (n) | Signed-w Spearman | \|w\| Spearman |
|---|---|---|---|---|---|
| s42 | Hill | 0.531 | 0.4856 (341) | 0.1901 | −0.0054 |
| s43 | Hill | 0.528 | 0.4867 (332) | 0.2393 | −0.0186 |
| s44 (dt0.25) | Hill | 0.534 | 0.4831 (347) | 0.1648 | −0.0275 |
| basin | Hill | 0.531 | 0.4856 (341) | 0.1901 | −0.0054 |
| prior | Hill | 0.606 | 0.5217 (362) | 0.2297 | 0.0588 |
| blindv2 | Hill | 0.558 | 0.5607 (362) | 0.3414 | 0.1026 |
| blind3 | tanh | 0.558 | 0.5509 (361) | 0.3348 | 0.0822 |
| blind4 | tanh | 0.558 | 0.5771 (362) | 0.2601 | 0.1244 |
| prior3 | tanh | 0.533 | 0.5369 (362) | 0.2950 | 0.0585 |
| prior4 | tanh | 0.612 | 0.5695 (362) | 0.2283 | 0.1215 |

**Pooled bands: Hill [0.528, 0.606] / tanh [0.533, 0.612]**

**Layer-level extrema** (different scale; not comparable to pooled bands): [0.387, 0.577]

Note: Pooled values are tie-averaged AUROC over the full candidate domain per arm; the layer-level extrema report per-layer minima/maxima across arms and are not directly comparable to the pooled bands; basin reuses the seed-42 run (s42) and is not an independent replicate. GE-domain AUROC is a recomputation over the GE candidate edges scored for each arm (n in parentheses) and is not directly comparable with pooled AUROC over the full six-class candidate domain. Signed-w and |w| Spearman columns report signed-weight and absolute-weight correlations, respectively.

blindv2: second execution of the blind-initialization arm (Hill family); the first attempt (initialization amplitude 0.4) stalled from epoch 1 and was abandoned, and the re-run at amplitude 0.05 trained successfully.

The basin row is the Hill-family s42 run scored for rank-level AUROC only; it is not the tanh matched-form basin curve of §2.4.

#### Supplementary Table S3. Rat five-layer differential analysis hits

| Layer | Features tested | Group hits | Sex hits |
|---|---|---|---|
| transcript | 21069 | 33 | 8361 |
| protein | 75229 | 1 | 1920 |
| phospho | 25660 | 0 | 635 |
| acetyl | 6180 | 21 | 1474 |
| metabolite | 554 | 100 | 361 |

**Note: Primary contrast (8-week completers vs remaining 36 animals); sex as the sole covariate; Benjamini-Hochberg adjusted $p < 0.05$ within each layer; this analysis serves as a framework-transfer check, not a mechanistic claim about rat exercise biology.**

#### Table S4. dynGENIE3 run-level scores (30 runs)

Run-level tie-averaged AUROC within the GE candidate domain (228 of 239 GE-layer genes observed under both conditions).

Two observation modes over the same ground-truth universe × three tree settings × five seeds. Raw dynGENIE3 output: VIM[regulator, target]; scoring used its transpose, VIM^T[target, regulator], with both index directions checked.

| Observation mode | Trees | Seed | AUROC |
|---|---|---|---|
| sim2i (ideal-data) | 100 | 0 | 0.555 |
| sim2i | 100 | 1 | 0.5732 |
| sim2i | 100 | 2 | 0.5279 |
| sim2i | 100 | 3 | 0.5592 |
| sim2i | 100 | 4 | 0.5422 |
| sim2i | 500 | 0 | 0.6198 |
| sim2i | 500 | 1 | 0.608 |
| sim2i | 500 | 2 | 0.5942 |
| sim2i | 500 | 3 | 0.5914 |
| sim2i | 500 | 4 | 0.5767 |
| sim2i | 1000 | 0 | 0.6124 |
| sim2i | 1000 | 1 | 0.6073 |
| sim2i | 1000 | 2 | 0.611 |
| sim2i | 1000 | 3 | 0.6195 |
| sim2i | 1000 | 4 | 0.5858 |
| sim2 | 100 | 0 | 0.6093 |
| sim2 | 100 | 1 | 0.4734 |
| sim2 | 100 | 2 | 0.5432 |
| sim2 | 100 | 3 | 0.4881 |
| sim2 | 100 | 4 | 0.5731 |
| sim2 | 500 | 0 | 0.5463 |
| sim2 | 500 | 1 | 0.5423 |
| sim2 | 500 | 2 | 0.5397 |
| sim2 | 500 | 3 | 0.5657 |
| sim2 | 500 | 4 | 0.5759 |
| sim2 | 1000 | 0 | 0.5161 |
| sim2 | 1000 | 1 | 0.5842 |
| sim2 | 1000 | 2 | 0.5897 |
| sim2 | 1000 | 3 | 0.5372 |
| sim2 | 1000 | 4 | 0.583 |

**Summary (registered values, §4.7): sim2i 1000-tree tier 0.607 ± 0.016; sim2 1000-tree tier 0.562.**

For the sim2i 1000-tree summary, 0.607 ± 0.016 denotes the mean ± 95% t-based confidence interval across five seeds.

### Supplementary Note S1. Rat cohort sensitivity analyses

The sensitivity set comprised 18 animals — 9 that completed 8 weeks of endurance training and 9 eight-week controls (10 females, 8 males) — drawn from the 45-animal analysis set defined in Methods (fixed before analysis).

**Table N1a. Covariate-adjusted differential analysis (sensitivity primary within supplement)**

| Layer | Features tested (BH denominator) | Group effect adj_p < 0.05 | Sex effect adj_p < 0.05 |
|---|---|---|---|
| transcript | 19970 | 3 | 5110 |
| protein | 75254 | 0 | 152 |
| phospho | 25711 | 0 | 24 |
| acetyl | 6184 | 0 | 390 |
| metabolite | 554 | 4 | 281 |

### Table N1b. Sex-stratified results (transparency listing)

| Layer | min raw p (F) | min raw p (M) |
|---|---|---|
| transcript | 2.28e-05 | 2.73e-05 |
| protein | 1.32e-04 | 1.11e-04 |
| phospho | 6.64e-03 | 3.43e-04 |
| acetyl | 6.19e-04 | 1.11e-04 |
| metabolite | 9.14e-04 | 5.26e-04 |

### N1b note: stratified p values are raw (uncorrected); no multiple-testing correction was applied, consistent with the transparency-only (no hypothesis testing) framing of the stratified listing.

Covariate-adjusted primary; stratified listed for transparency, no hypothesis testing.

Sensitivity analysis restricted to 8-week trained versus 8-week control animals (n = 9 each), accompanying the primary eight-week contrast in the main text; adjusting for sex.

Given the small per-arm sample, per-sex subgroups (approximately n = 4-5 per cell) are shown for completeness only; no subgroup-level hypothesis testing is intended.
